# Real-Time Assessment of Murine Cardiac Oxygenation Using Photoacoustic Imaging

**DOI:** 10.64898/2026.06.30.735674

**Authors:** Jennie Vu, Ibrahim Khodabocus, Simone Derzi, Matthew Henry, Sandra T. Davidge, Kimberly F. Macala, Stephane L. Bourque, Ronan Noble

**Author notes:** Corresponding Author Ronan Noble, Department of Anesthesiology, Perioperative and Pain Medicine, University of Calgary, Calgary, Alberta, Canada, T2N 4N1.

## Abstract

**Background:** Perioperative incidents such as hypoxic cardiac injury often have subtle or nonspecific clinical manifestations. Reduction in myocardial oxygenation precedes biochemical changes, as well as electrical and functional changes. Photoacoustic imaging (PAI) is a modality that uses laser irradiation of tissue to generate ultrasonic waves, enabling spatially resolved quantitative mapping of oxygenated and deoxygenated haemoglobin. We investigated the utility of PAI for real-time monitoring of myocardial and great vessel oxygenation.

**Methods:** Male CD-1 mice were anaesthetised, and photoacoustic and simultaneous B-mode images were acquired of the myocardium and right ventricular outflow tract (RVOT), the pulmonary artery, and aorta. PAI was performed at fractional inspired oxygen levels (FiO_2_) of 100%, 21%, and then 10%. Separate cohorts of mice were exposed to increasing intravenous doses of either combined phenylephrine and isoprenaline, or individual administration of vasoactive or adrenergic agents.

**Results:** PAI reliably distinguished changes in oxygenation in the RVOT cavity, pulmonary artery, aorta, and myocardium. PAI detected hypoxia-induced changes in oxygenation, revealing greater desaturation in the myocardium than in the RVOT (−9.85%, 95% CI -14.94 to -4.77, P<0.0001). Escalating doses of phenylephrine and isoprenaline caused a progressive desaturation of the myocardium and RVOT (mean [95% CI]; myocardium 16 mg/kg: -14.64% [-27.62 to -1.65], P=0.0038 and RVOT 32 mg/kg: -18.71% [-32.15 to -5.27], P=0.0003). Myocardial deoxygenation was detected before changes in systolic function or electrical abnormalities.

**Conclusions:** This work demonstrates that PAI can reliably monitor cardiac oxygen desaturation, potentially offering an earlier warning of cardiac dysfunction and injury compared to existing monitoring tools.

## INTRODUCTION

The myocardium is highly susceptible to oxygen supply-demand mismatch, particularly in the perioperative setting where physiological stress is common. In fact, perioperative myocardial injury is predominantly caused by oxygen supply-demand imbalance; this imbalance may result in acute and clinically silent episodes of myocardial deoxygenation, contributing to myocardial injury and subsequent morbidity and mortality ^1–3^.

However, these events often go undetected, as current monitoring modalities such as changes in blood pressure, heart rate, urine output, capnography or the presence of cyanosis (primarily reflecting physiological consequences rather than imminent events) are insufficiently sensitive or rapid to capture early deterioration. Similarly, laboratory findings including base excess, lactate, and mixed venous oxygen saturation, together with electrocardiographic changes may be subtle, nonspecific, or lacking ^4, 5^. Although invasive monitors such as central venous and pulmonary artery catheters may provide information, their use is limited by cost, procedural complexity, and associated risk ^6^. Definitive imaging modalities such as coronary angiogram is generally impractical in perioperative settings or in critically ill patients. Consequently, clinically significant haemodynamic insults such as oxygen supply-demand mismatch leading to myocardial injury are often undetected, thereby delaying timely intervention that could reduce morbidity and mortality, ultimately contributing to irreversible damage and clinical deterioration ^7^.

Cardiac ultrasound is the most commonly used modality for haemodynamic assessment in the intensive care unit and emergency department ^8, 9^, and is similarly employed in the operating room by anaesthesiologists for managing unstable patients and major surgical cases via transthoracic or transoesophageal approaches ^1, 10–13^. In the context of myocardial assessments, reductions in oxygenation and perfusion precede biochemical changes, as well as changes in ECG and wall motion abnormalities ^14^. Given that ECG and ultrasound do not directly measure oxygenation, tissue damage therefore has the potential to occur due to delay in detection. In fact, Blood Oxygen Level Dependent Magnetic Resonance Imaging (BOLD MRI) has been shown to detect myocardial oxygen changes prior to or even in the absence of ECG and coronary perfusion changes, but it is not accessible intraoperatively or at the bedside ^15, 16^. While ultrasound remains an indispensable clinical tool, no existing imaging modality offers real-time, spatially resolved monitoring of cardiac oxygenation within the operating room or at the bedside.

Photoacoustic imaging (PAI) uses laser irradiation of a tissue to generate ultrasonic waves that can then be detected by an ultrasound transducer. When overlaid with ultrasound images, PAI signals enable real-time and spatially resolved quantitative mapping of various molecules (e.g. oxygenated and deoxygenated haemoglobin) based on absorption spectra ^17, 18^. Unlike pulse oximetry, which is limited to the periphery and thus may reflect delayed changes ^19–22^, PAI can resolve focal regions of interest in real-time. Indeed, while PAI has primarily been applied to study tumor growth and vascularization ^23^, our group has recently used this technology to study the oxygenation of the fetoplacental unit ^24^. However, if able to monitor oxygenation within the great vessels as well as the myocardium (by measuring the oxygenation within coronary microcirculation ^25^) in real-time, PAI would be positioned uniquely amongst cardiovascular imaging modalities. Its potential advantages in clinical anaesthesia include identifying when surgical stress may be poorly tolerated in a critically ill patient or in those at greater risk of myocardial injury (e.g. cardiovascular comorbidities, major surgeries), as well as providing new physiological targets to which interventions may be more closely titrated (e.g. medications, blood products, ventilation, and oxygenation).

The primary objective of this study was to evaluate the application of PAI in monitoring oxygenation of the myocardium and great vessels and benchmark its performance against conventional ECG and echocardiography as a potential early warning sign for myocardial hypoxaemia in mice. As PAI provides a direct measurement of oxygenation, we hypothesized that measurable changes would present prior to changes in myocardial contractility or conduction. We then sought to test the murine myocardial response to select vasoactive and inotropic agents commonly found in the operating room, ICU, and CCU.

## METHODS

### Animal model

All animal procedures were reviewed and authorized by the University of Alberta Animal Care Committee (protocol #3881) and adhered to the guidelines established by the Canadian Council on Animal Care and the National Institutes of Health Guide for the Care and Use of Laboratory Animals. Naïve male CD-1 mice (10-12 weeks old, 45-50 g, Charles River, Kingston, NY) were anaesthetized with isoflurane (4% induction, 2% maintenance) carried in 1 L/min medical-grade gas (composition described below in protocols 1 through 4) and placed on a heated ultrasound platform in a supine position. The ventral thoracic region was depilated (Nair®) for 30 seconds (s). Adequate surgical anaesthesia was confirmed by absence of the limb withdrawal reflex to toe pinch prior to instrumentation; continuous heart rate monitoring was performed, and bradycardia below 100 bpm prompted humane euthanasia.

### Photoacoustic and ultrasound imaging

Imaging experiments were performed using the Vevo 3100 ultrasound system in conjunction with an integrated LAZR-X photoacoustic platform (VisualSonics Inc., Toronto, Canada) using an optical fibre width of 24 mm paired with the MX400 (20-46 MHz) ultrasound probe, resulting in an axial resolution of 50 μm and temporal resolution of 0.2 s.

Prior to imaging, the laser was calibrated with an external energy sensor, and limb leads were secured to confirm a stable three lead ECG. For all myocardial assessments, B-mode and photoacoustic (PA)-mode images were acquired in the parasternal long-axis view, which positioned the anteroseptal wall of the left ventricle toward the most superficial aspect of the thorax. In pilot experiments, this orientation maximized sensitivity for assessing systolic function (fractional area change and longitudinal strain) whilst allowing photoacoustic measurements of the full longitudinal extent of the anterior myocardium. Imaging was repeated using a modified, basally oriented parasternal short-axis view to measure photoacoustic signals from the pulmonary artery and aorta. For each image, the heart or great vessels were positioned within the optical focus between 10 and 17 mm, ensuring the field was not obstructed by artifacts (e.g. bubbles, hair, skin reverberations). Time gain compensation was adjusted to enhance deeper structures and was maintained consistently across animals. The images used to quantify oxygen saturation were collected using 750 and 850 nm wavelengths in the built-in Oxy-Hemo mode.

### Hypoxia-reoxygenation

First, to externally validate PAI arterial measurements, mice were anaesthetized as described above with a fractional inspired oxygen (FiO_2_) of 100% (medical grade oxygen), and a standard pulse oximeter was placed on the left hind paw (PhysioSuite^TM^, Kent Scientific, Torrington, CT). Next, four imaging protocols were conducted in three separate cohorts of mice.

Protocol 1 (Cohort 1): Baseline photoacoustic and simultaneous B-mode ultrasound images of the myocardium microvasculature and blood within the right ventricular outflow tract (RVOT), as well as the pulmonary artery and aorta were initially acquired at steady state with a FiO_2_ of 21% (medical grade air, 79% nitrogen) and then 100% (medical grade oxygen) (n=5-7). Within each imaging window, we compared adjacent arterial and venous counterpart structures as internal references at the same acoustic depth; accordingly, myocardial tissue was evaluated alongside the RVOT in the long-axis view, and the aorta was compared with the pulmonary artery in the short-axis view. Arterial and venous sO_2_ values were compared to assess signal fidelity. Subsequent analyses focused on the myocardium and RVOT due to the clinical importance of myocardial oxygenation and the larger analysable surface area afforded by this imaging plane. Following each assessment, a second image was taken using the electrocardiogram (ECG)-gated Kilohertz Visualisation (EKV) mode, which is a high-frame-rate ultrasound technique that visualizes rapid cardiac motion in small animals by averaging several cardiac cycles recorded at 250 Hz to facilitate ECG-gated reconstruction of a representative cardiac cycle. This was piloted prior to continuous recordings, to assess how cardiac and respiratory motion affected imaging quality.

Protocol 2 (Cohort 1): In the same cohort of mice, steady state, equilibrated recordings were acquired of the myocardium and RVOT at a FiO_2_ of 100%. The FiO_2_ was then reduced to 21% by changing gas delivery to 100% medical grade air and the response was recorded for 4 min. FiO_2_ was then reduced to 10% by introducing a 50:50 mix of medical grade air and nitrogen through a gas mixer for 60 s. FiO_2_ was then restored to 100% for 120 s to capture the reoxygenation response (n=11). Finally, FiO_2_ was reduced to 0% to observe lowest physiological PAI signals within the myocardium. System clock time and frame numbers were recorded during acquisition to facilitate downstream analyses.

To assess large vessel oxygenation dynamics, the same sequence of FiO_2_ changes (100% → 21% → 10% → 100%) was performed whilst imaging the pulmonary artery and aorta (n=4). Mice were concurrently instrumented with a pulse oximeter (PhysioSuite, Kent Scientific) placed on the hind paw for SpO_2_ measurements. Pulse oximetry recording timestamps were synchronised with the photoacoustic system to enable direct comparison of values.

### Photoacoustic monitoring of cardiac supply-demand mismatch

Protocol 3 (Cohort 2): CD-1 mice were anaesthetized as described above (FiO_2_ of 100%), and ventrally depilated. Adequate depth of anaesthesia was confirmed by absence of withdrawal to hind paw pinch prior to femoral vein cannulation. Mice were thereby instrumented with a femoral vein cannula (26G) for intravenous infusions (n=8); experimental setup depicted in **Suppl Figure S1**. Baseline PA imaging (5 min.) of the myocardium was performed with simultaneous B-mode ultrasound acquisition, with a concerted effort to capture clear recordings of the anteroseptal myocardial wall - a region conducive to oxygenation assessments by PAI. Mice received an IV bolus of phenylephrine (17205, Cayman Chemicals) and isoprenaline (I6504, Sigma Aldrich) at a 2:1 ratio (with concentrations of 10 mg/mL each, combined into a single syringe) to model increased cardiac metabolic demand by simultaneously increasing cardiac contractility (via β stimulation) and increasing afterload (via α stimulation). Boluses were doubled every 8 minutes, with 8-minute recording periods between doses (volumes of 20 → 320 µL, corresponding to doses of 2.66 → 42.67 mg/kg for phenylephrine, and 1.33 → 21.33 mg/kg for isoprenaline), until myocardial oxygen saturation decreased by ∼40% from baseline. Combined dosages of 4 → 64 mg/kg are reported in figures. Animals failing to reach a 40% decrease in sO_2_ were excluded from supply-demand mismatch analyses; two mice were excluded.

Protocol 4 (Cohort 3): In a third cohort of mice, the effects of increasing doses of phenylephrine, isoprenaline, and noradrenaline, and sodium nitroprusside in isolation were assessed (n=3 each). Each mouse was randomly assigned to be administered one of the four drugs in increasing doses of: 4 to 16 mg/kg phenylephrine; 4 to 32 mg/kg isoprenaline; 4 to 16 mg/kg sodium nitroprusside (PHR1423, Sigma Aldrich); 0.4 to 12.8 mg/kg noradrenaline (A7257, Sigma Aldrich) until myocardial oxygen saturation decreased by 40%.

For all cohorts, VevoLab 5.8.0 was used to perform spectral unmixing on images acquired in photoacoustic (PA)-mode; regions of interest were demarcated based on structures identified within B-mode images to retrieve measurements of average oxygen saturation signal (oxy-haemoglobin signal divided by combined oxy- and deoxy- haemoglobin amounts to generate a percentage, sO_2_) and average total haemoglobin (HbT, reflecting myocardial perfusion) normalized to the outlined region area. Photoacoustic and functional analyses (to determine regional sO_2_ and systolic parameters) were conducted by an operator blinded to drug identity but unblinded to treatment timing. Time from intervention to first significant deviation was compared across modalities. Functional analyses were restricted to mice with stable baseline ECG signals, ultrasound B-mode frames with clearly defined endocardial borders, delineation of the myocardial wall, and aortic valve visualization to confirm the correct imaging plane.

### *In vivo* haemodynamic characterisation

Protocol 5 (Cohort 4): Central arterial blood pressure was measured *in vivo* using a 1.0F pressure–volume catheter (Millar, ADInstruments) interfaced with a PowerLab/MPVS Ultra system (ADInstruments). Mice were anaesthetised (see above) and a surgical plane of anaesthesia confirmed by absence of limb withdrawal reflex. The right common carotid artery was isolated and an arterial line was placed; the catheter was thereby advanced to a region above the aortic arch. The jugular vein was isolated for cannulation. After a 5-min baseline recording, mice received sequential intravenous phenylephrine–isoprenaline dosing (2.66 → 42.67 mg/kg for phenylephrine, and 1.33 → 21.33 mg/kg for isoprenaline), with 8-min recording intervals between doses (n=6). Individual drugs were administered in increasing doses: 4 to 16 mg/kg phenylephrine; 4 to 32 mg/kg isoprenaline; 4 to 16 mg/kg sodium nitroprusside; 0.4 to 12.8 mg/kg noradrenaline (n=3 each). Haemodynamic data was analysed using LabChart 8 software. Echocardiography was performed at 2-minute intervals using a Vevo 2100 system, with resultant data analysed using VevoLab 5.8.0 software.

### ECG analyses

Lead II electrocardiogram profiles were acquired using the Vevo 3100 platform after limbs were secured to four limb leads (RA, LA, LL, RL) with electrode gel and surgical tape. ECG abnormalities were identified as alterations in QRS complex, J-wave duration and amplitude (a sign of myocardial hypoxia in mice) ^26, 27^, appearance of bundle branch block conduction patterns, and prolongation of PR and QT intervals.

### Plasma cardiac Troponin T and Lactate measurements

After imaging (∼45 min total duration) mice were exsanguinated under anaesthesia by carotid-cut, and whole blood was collected in heparin-lined tubes. Blood was also collected from age, sex and weight-matched naïve control mice that underwent the same instrumentation and imaging protocols but were not exposed to FiO_2_ changes and did not receive intravenous pharmacological agents. Blood samples were centrifuged for 15 min at 3500×G at 4°C. The resulting plasma was then aliquoted and snap frozen in liquid nitrogen. Plasma samples underwent one freeze-thaw cycle for measurements of cardiac Troponin T (cTnT) using a Mouse Cardiac Troponin T ELISA Kit (Antibodies, A73988, detection range 9.37-1000 pg/mL) according to the manufacturer’s instructions. Blood lactate was measured using a handheld analyser (Lactate Plus Meter, Nova Biomedical). A baseline value was obtained from a small-volume blood sample collected before physiological recordings. At the end of the protocol, lactate was measured from whole blood collected by carotid exsanguination. Blood was collected only from animals with beating hearts at the time of sampling.

### Statistical Analyses

All n values reflect biological replicates. We performed power calculations based on myocardial desaturation (sO_2_ reduction), the primary outcome measure and most variable endpoint, to guide study design. Assuming a two-tailed α of 0.05 and power (1–β) of 0.8, the minimum sample size was 3 mice per group for individual drug studies and 5 mice for hypoxia and combination challenges. Data were analysed in GraphPad Prism 11 by two-way ANOVA for steady state data, or by fitting a repeated-measures two-way ANOVA mixed-effects model for the effects of region (i.e. myocardium vs right ventricular outflow tract) and time. Data were assessed for normality and homogeneity of variance prior to one-way and two-way ANOVA analyses and were transformed if heteroscedastic. If assumptions were met, standard one-way or two-way ANOVA was performed, and non-normally distributed data were analysed using the Kruskal-Wallis test. For pairwise comparisons, normally distributed data were analysed using Student’s t-test, whereas non-normally distributed data were analysed using the Mann-Whitney U test, while a paired t test was used for blood lactate measurements. Tukey’s or Sidak’s post-hoc tests were used for multiple comparisons following two-way ANOVAs, whilst Dunnett’s post hoc test was used following one-way ANOVAs.

## RESULTS

### Baseline Photoacoustic Imaging of Myocardial and Great Vessel Oxygenation

In the first protocol, oxygen saturation (sO_2_) was measured in four regions at two fractions of inspired oxygen (FiO_2_) conditions. Comparisons were made between the anteroseptal myocardium and RVOT (**Figure 1A, Figure S2A**) and great vessels (**Figure 1B**) to assess PAIs ability to distinguish regional differences in sO_2_ in real time under varying oxygenation statuses. PAI reliably distinguished sO_2_ in the RVOT cavity, pulmonary artery (Pulm Art), aorta, and anteroseptal myocardium presented as mean [95% CI]: 75.8% [72.7-78.94], 78.0% [73.3-82.8], 86.6% [83.8-89.4], 86.3% [83.7-89.0], and 67.4% [61.0-73.7], 69.2% [63.6-74.8], 77.0% [71.1-83.0], 72.5% [66.5-78.6] at FiO_2_ of 100% and 21% respectively (**Figure 1C, D**). Time gain compensation aided effective measurement of consistent sO_2_ values between the RVOT and Pulm Art while preserving detectable differences due to FiO_2_ changes, despite data acquisition at different thoracic depths (**Figure S2C**). Standard PAI showed that at FiO_2_ 100%, but not 21%, RVOT signals were lower than myocardial sO_2_ values (mean difference [95% CI]; FiO_2_ 100%: 10.51% [2.78–18.25], P=0.0046; FiO_2_ 21%: 5.16% [2.57 to 12.90], P=0.33), consistent with venous blood being comparatively deoxygenated relative to mixed blood sources (arterial and venous) within the myocardium (**Figure 1C**). Similarly, at each FiO_2_, pulmonary artery sO_2_ was lower than aortic sO_2_ (mean difference [95% CI]; FiO_2_ 100%: 8.59% [1.023-16.16], P=0.022; FiO_2_ 21%: 7.802 [0.2346-15.37], P=0.042) **(Figure 1D**). PAI-derived aortic sO_2_ showed strong agreement with pulse oximetry SpO_2_, with high Pearson’s R and Lin’s concordance correlation coefficient values of 0.979 and 0.971 respectively (**Figure 1G**). Bland-Altman analysis revealed agreement at physiological ranges of sO_2_, but increasing discordance at subphysiological sO_2_ values, where pulse oximetry occasionally failed to report values (SpO_2_ < 60%) (**Figure 1H**). PAI showed modest positive bias at low sO_2_ and negative bias at higher sO_2_ conditions, with an overall mean difference of ∼2.5% across FiO_2_ conditions with 95% limits of agreement of -7.82 to 3.32 and -6.89 to 3.90 for FiO_2_s of 100 and 21% respectively (**Figure 1H-I**).

**Figure 1.**
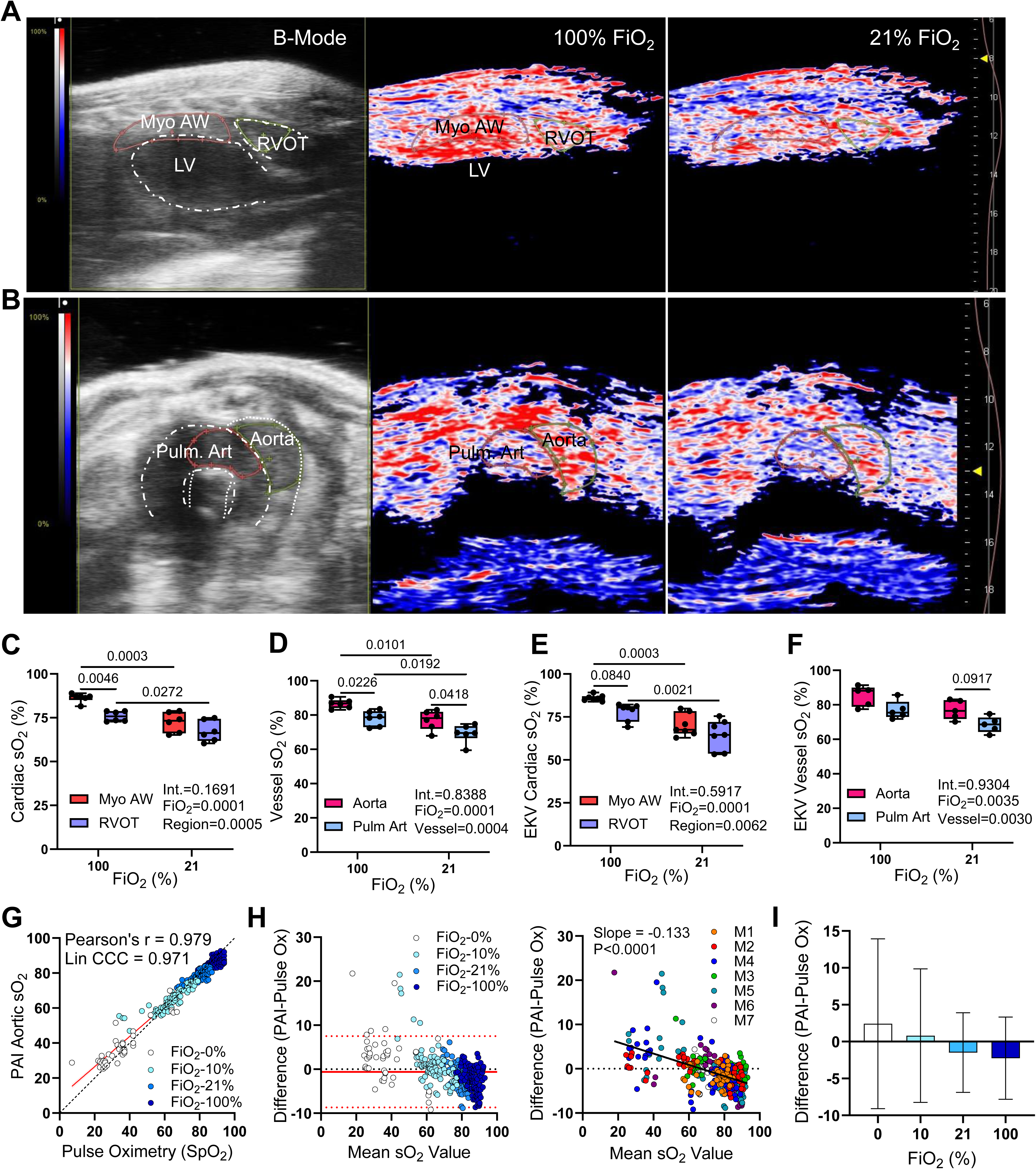
Photoacoustic imaging of cardiac and great vessel oxygenation. (**A**) Parasternal long-axis B-mode ultrasound identifying regions of interest (ROIs) for photoacoustic imaging (PAI). The anteroseptal myocardial wall is outlined in pink, the LV cavity is outlined in dashed white, and the right ventricular outflow tract (RVOT) is outlined in green and dashed white (left panel). Corresponding PAI images acquired at 100% FiO_2_ (middle panel) and 21% FiO_2_ (right panel) are shown, where bright red indicates 100% oxygen saturation (sO_2_) and dark blue/black indicates 0% sO_2_. (**B**) Modified parasternal short-axis B-mode ultrasound depicting ROIs of the great vessels, with the aortic arch outlined in green (demarcated with a dashed white line) and the main pulmonary artery outlined in red (demarcated with an interrupted white line) (left panel). Corresponding PAI images acquired at steady state under 100% FiO_2_ (middle panel) and 21% FiO_2_ (right panel) are shown. Bright red indicates 100% sO_2_, dark blue/black indicates 0% sO_2_. Skin-line artifacts are visible at the inferior aspect of the image outside the ROIs. (**C**) Quantification of regional sO_2_ in the anterior myocardial wall (Myo AW) and right ventricular outflow tract (RVOT) in the standard photoacoustic imaging mode, n=6. (**D**) Quantification of regional sO_2_ in the great vessels, including the aorta and pulmonary artery (Pulm Art) in the standard photoacoustic imaging mode, n=6. (**E**) ECG-gated kilohertz acquisition of regional sO_2_ dynamics with corresponding quantification of Myo AW and RVOT oxygenation, n=7. (**F**) ECG-gated kilohertz acquisition of regional sO_2_ dynamics with corresponding quantification of aortic and pulmonary artery oxygenation, n=5. (**G**) Correlation of pulse-oximetry derived SpO_2_ (x axis) and PAI-derived SaO_2_ (y-axis) across FiO_2_ conditions (0, 0.1, 0.21, 1.00). Lin’s Concordance Correlation Coefficient, Lin CCC; Pearson’s r, Pearson correlation coefficient, n=7. (**H**) Bland-Altman plots showing agreement between PAI SaO_2_ and pulse oximetry SpO_2_. The mean difference (bias, solid red line) and 95% limits of agreement (dashed red lines) are indicated. Each point represents an individual technical replicate measurement, plotted as the difference versus the mean of the two methods. The left plot clusters individual values based on FiO_2_, while the right plot clusters values based on biological replicates, n=7 (**I**) Absolute biases derived from Bland-Altman analyses at each FiO_2_ condition, with 95% limits of agreement indicated, n=7. Int., interaction effect; FiO_2_, differences in fractional inspired oxygen levels; Region, denotes differences in oxygen saturation between anatomical regions; Vessel, denotes differences in oxygen saturation between great vessels; Data are expressed as mean and IQR, with minimum and maximum values. Comparisons between groups were performed using two-way ANOVA followed by Tukey’s post hoc test.

Relative to standard PAI, motion-gated EKV-acquired steady-state measurements were less sensitive in detecting sO_2_ differences between the great vessels (aorta and pulmonary artery) and similarly or less able to measure sO_2_ changes across the different FiO_2_ levels (**Figure 1E,F**). Motion-gated imaging (**Figure 1E, F**) requires 90 to 300 s to capture a single averaged cardiac cycle, leading to challenges when measuring changes over prolonged periods. No discernible differences in myocardial sO_2_ were observed between systole and diastole using either EKV or standard photoacoustic imaging despite EKV’s motion-gated design (**Figure S2D**). Both modalities demonstrated comparable variance (standard deviation and coefficients of variation) during steady-state baseline myocardial sO_2_ measurements (**Figure S2E**). Given that EKV mode was not superior to standard photoacoustic imaging for distinguishing cardiac sO_2_ values and limited the length of recordings, we proceeded with continuous acquisitions without cardiac motion correction.

### Myocardial and Great Vessel Oxygenation Across Dynamic Hypoxic Ranges

In the second protocol, continuous image recordings were performed first of the myocardium and RVOT, and then of the aorta and pulmonary artery (**Figure S2A**). When FiO_2_ was reduced from 100% to 21%, myocardial and RVOT sO_2_ decreased (mean [95% CI]; Myo AW: -9.15% [-11.57 to -6.74] and RVOT: -8.53% [-11.93 to -5.122] respectively, P<0.0001, **Figure 2C**), as did aortic and pulmonary artery saturations (mean [95% CI]; Aorta: -12.15% [-19.54 to -4.77] and Pulm Art: -10.2% [-15.42 to -4.99] respectively, P<0.0001 **Figure 2D**), though regional differences in sO_2_ persisted between Myo AW and RVOT (**Figure 2C, Suppl Figure S3A**), but not between aorta and pulmonary artery (**Figure 2D, Suppl Figure S3B**). There were no differences in either absolute or relative desaturation between the Myo AW and the RVOT (**Figure 2C, Suppl Figure S3A, S4C**), or between the aorta and pulmonary artery (**Figure 2D, Suppl Figure S3B, S2B, S4D**).

**Figure 2.**
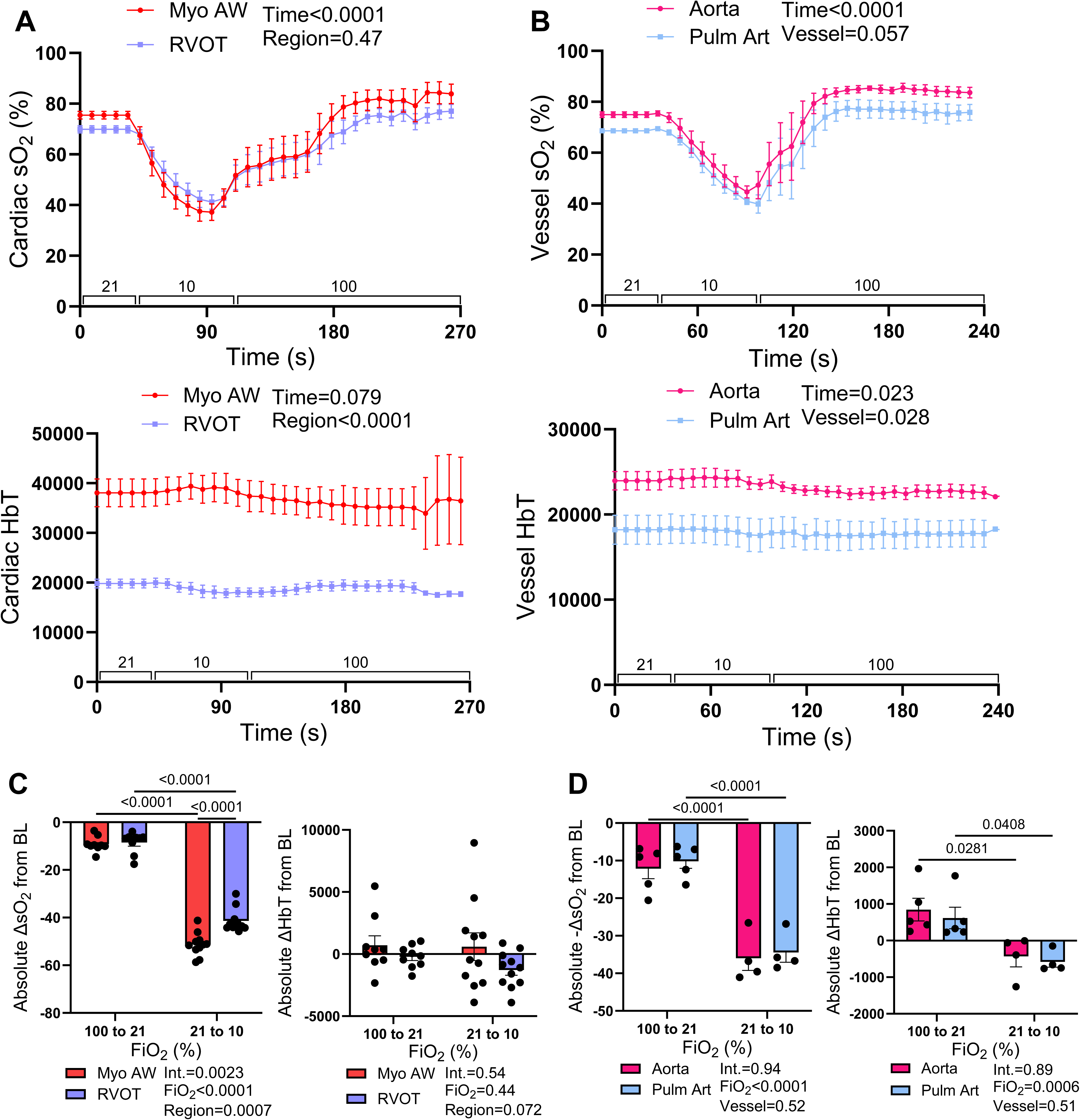
Photoacoustic measurements of regional oxygenation during acute hypoxia. (**a**) Oxygen saturation (sO_2_, above) and total haemoglobin concentration (HbT, below) over time from FiO_2_s of 21% (baseline) to 10%, and 100%, of the myocardial anterior wall (Myo AW) and right ventricular outflow tract (RVOT), n=11. (**b**) Oxygen saturation (above) and haemoglobin concentration (below) over time from FiO_2_s of 21% (baseline) to 10%, and 100% of the aorta and pulmonary artery (Pulm Art), n=4. (**c**) Absolute changes in oxygenation (left) and haemoglobin concentration (right) from baseline in the Myo AW and RVOT in response to FiO_2_ transitions of 100% to 21% and 21% to 10%, n=11 (**d**) Absolute changes in oxygenation (left) and haemoglobin concentration (right) from baseline in the aorta and pulmonary artery in response to FiO_2_ transitions of 100% to 21% and 21% to 10%, n=4. Data are expressed as mean±SEM. Comparisons between groups were performed using either a mixed-effects repeated measures 2-way ANOVA followed by Tukey’s post-hoc analysis or a one-way ANOVA followed by the Dunnett’s post hoc test.

When FiO_2_ was reduced from 21% to 10%, sO_2_ of the myocardium and RVOT decreased by - 51.23%, 95% CI -54.50 to -47.95 and -41.37%, 95% CI -44.62 to -38.13, P<0.0001 respectively after 90 s (at peak hypoxia; **Figure 2A,C; Suppl Figure S4C**). Similarly, aortic and pulmonary artery sO_2_ decreased by -35.98%, 95% CI -46.37 to -25.59 and -34.46%, 95% CI -42.71 to - 26.20, P<0.0001 respectively at 90 s after FiO_2_ 10% (peak hypoxia; **Figure 2B,D; Suppl Figure S4D**). The Myo AW exhibited a greater absolute and relative reduction in sO_2_ compared with the RVOT in response to FiO_2_ 10% (mean -9.85%, 95% CI -14.94 to -4.77, p<0.0001, **Figure 2C**, **Suppl Figure S9A**). These differences were not attributable to alterations in blood flow, as reductions in total haemoglobin (HbT) did not differ between regions (**Figure 2A, Suppl Figure S4A,C**). Pulm Art and aortic sO_2_ decreased to a similar degree throughout the FiO_2_ 10% challenge, with aortic sO_2_ values consistently exceeding Pulm Art values (**Figure 2B, D; Suppl Figure S4B, S9B**). Total haemoglobin (HbT, representing myocardial blood flow) was modulated by FiO_2_ alterations over time (**Figure 2A-D, Suppl Figure S3A-B**), though statistically significant or trending, the magnitude of these changes across time and between FiO_2_ challenges were physiologically modest. In both protocol one and two, HbT was recorded highest in the myocardium, and then the aorta, RVOT, and pulmonary artery respectively. To assess the lowest physiologically attainable sO_2_ measurements, FiO_2_ was transitioned from 21% to 0%. Oxygen saturation values in the Myo AW, RVOT, aorta, and pulmonary artery declined and plateaued at approximately 25–30%, likely reflecting residual oxygen occupancy of haemoglobin (**Suppl Figure S3C-D**).

### Pharmacologic induction of myocardial supply-demand mismatch

In the third protocol, we increased myocardial metabolic demand with combined phenylephrine and isoprenaline to model the increasing vasopressor/inotrope requirements of a critically ill patient (**Suppl Figure S1A-B, S2B**). We found that levels of plasma cTnT and blood lactate - markers of myocardial/tissue injury - were elevated (cTnT: mean difference [SEM], 47.5 pg/mL [18.7], P=0.0043, lactate: mean difference [95% CI]; 3.9 mmol/L, [1.79-6.01], P=0.0068) following titration of pharmacological agents compared to control and baseline animals (**Figure 3D**), consistent with clinical presentations of supply-demand mismatch (3).

**Figure 3.**
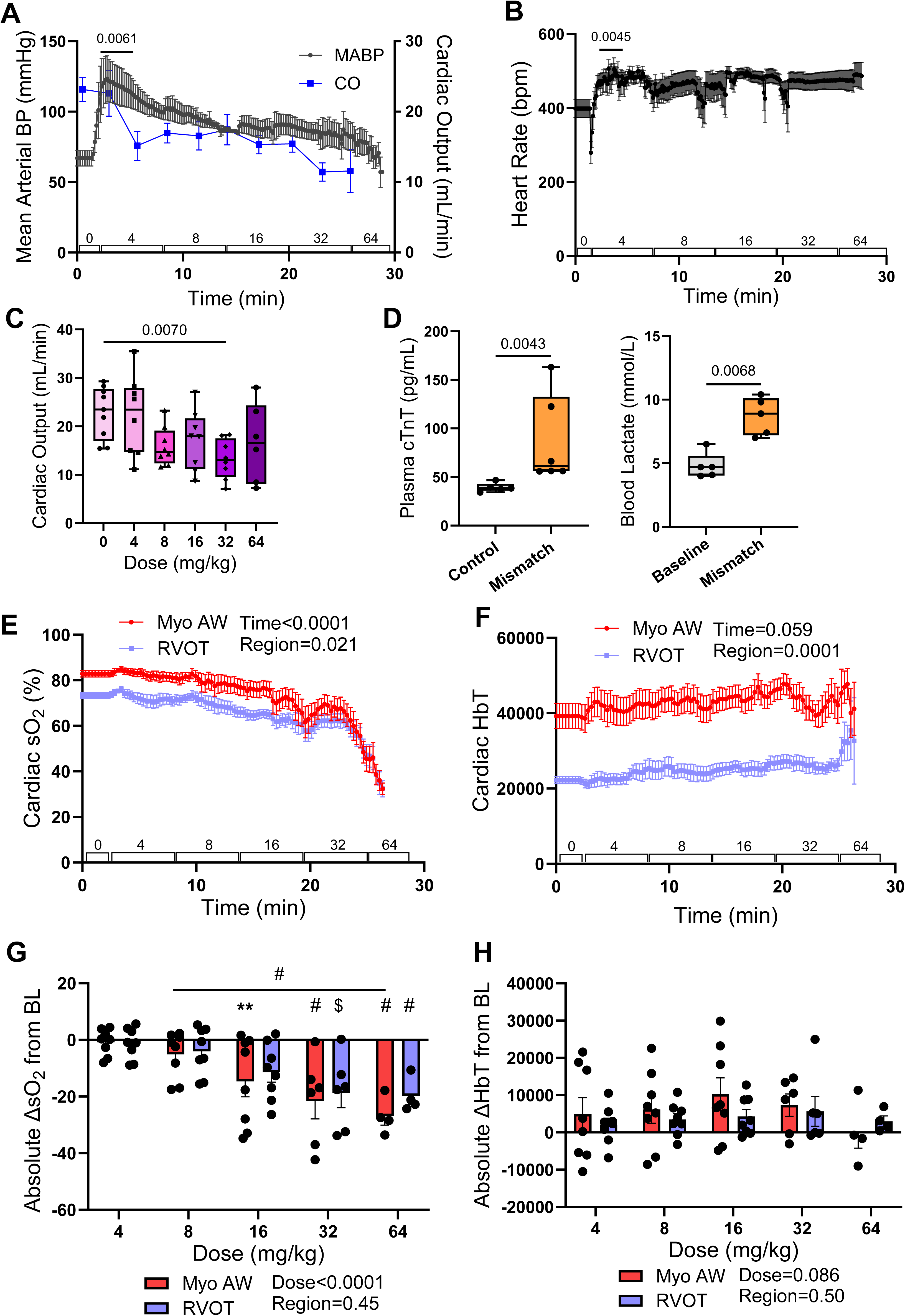
Photoacoustic imaging of cardiac oxygenation during pharmacologically induced supply-demand mismatch. (**A**) Mean arterial blood pressure (BP) measurements (left y-axis) from an indwelling carotid arterial line and cardiac output (blue line, right y-axis) measured across cocktail drug titration protocol, n=6. (**B**) Heart rate measured throughout the titration protocol. (**C**) Quantification of cardiac output at each combination (phenylephrine-isoprenaline) drug dose, n=6-8. (**D**) Plasma cardiac troponin T concentrations (left) and blood lactate (right) measured following completion of imaging and pharmacological challenge, n=5-6. (**E**) Oxygen saturation of the anteroseptal myocardium (Myo AW) and right ventricular outflow tract (RVOT) measured over time following baseline and sequential bolus administration of phenylephrine and isoprenaline delivered via a single syringe at 8-min intervals (20 → 40 → 80 → 160 → 320 → 640 µL), corresponding to cumulative doses of 2.66/1.33, 5.32/2.66, 10.64/5.32, 21.28/10.64, and 42.56/21.28 mg/kg of phenylephrine/isoprenaline, respectively. Total administered dose is indicated on the graph, n=8. (**F**) Haemoglobin concentration (HbT) measured in parallel across time during phenylephrine/isoprenaline titration. (**G**) Absolute changes in oxygenation of the Myo AW and RVOT compared to baseline in response to increasing dosages; statistical significance indicates comparison to dose 4 mg/kg, n=4-8. (**H**) Absolute changes in haemoglobin concentration in the Myo AW and RVOT compared to baseline in response to increasing dosages; statistical significance indicates comparison to dose 4 mg/kg, n=4-8. Data are expressed as mean±SEM, or mean and IQR, with minimum and maximum values. Comparisons between groups were performed using either a mixed-effects repeated measures 2-way ANOVA followed by Tukey’s post hoc test, or a one-way ANOVA followed by Dunnett’s multiple Comparisons. Mann-Whitney t test and paired t-test were used to compare cTnT and lactate respectively. *p* < 0.05, \*\**p* < 0.01, $*p* < 0.001, #*p* < 0.0001.

Induction of supply-demand imbalance was associated with an increase in mean arterial blood pressure by 53 mmHg, 95% CI 18.98 to 87.60 (**Figure 3A**, P_BP_ = 0.0061), accompanied by an approximate 94 bpm increase in HR (95% CI 34.25-153.8) from baseline (**Figure 3B**, P_HR_ = 0.0045) further highlighting the increased cardiac afterload and workload. Cardiac output was variable across doses but showed a downward trend, reaching a significant decrease only at 32 mg/kg of cocktail (**Figure 3C**). Induction of myocardial supply-demand mismatch also elicited a slight trend towards an enhanced myocardial blood flow, indicated by increasing total haemoglobin (HbT) measurements throughout the recording (**Figure 3F, H, Suppl Figure S5B**), consistent with metabolic upregulation of coronary flow ^28–30^. With escalating doses of the cocktail, we observed progressive myocardial desaturation as well as decreasing RVOT oxygenation (**Suppl Video 2**, **Figure 3E, G**). Although measurable baseline differences in sO_2_ were observed between the Myo AW and RVOT, equivalent levels of desaturation occurred at higher doses. Myo AW dose-specific significant sO_2_ reductions compared to 4 mg/kg sO_2_ presented as mean [95% CI]; 16 mg/kg: -14.64% [-27.62 to -1.651], P=0.0038; 32 mg/kg: -21.56% [-37.78 to -5.34], P<0.0001, and 64 mg/kg: -26.81% [-37.26 to -16.36], P<0.0001 and RVOT dose-dependent significant sO_2_ reductions of 32 mg/kg: -18.71% [-32.15 to -5.27], P=0.0003; 64 mg/kg: -19.71% [-29.54 to -9.87], P=0.0005 were observed (**Figure 3E, G**). Increased oxygen extraction in response to heightened metabolic demand likely underlies this desaturation ^31^. Absolute magnitudes of cardiac desaturation (compared to an initial dose of 4 mg/kg) reached significance at a total dose of 16 mg/kg but was most prominent following a total dose of 32 mg/kg and 64 mg/kg (21 and 43 mg/kg of phenylephrine and 11 and 21 mg/kg of isoprenaline) (**Figure 3G**, P<0.0001). Relative reductions in sO_2_ across time did not differ between the regions; overall relative trending increases in HbT (P_time_=0.059, P_dose_=0.086) over time were similar between the Myo AW and RVOT (**Suppl Figure S5A-D**). During this protocol, some instances of abrupt myocardial and RVOT deoxygenation were observed (**Suppl Video 1, Suppl Figure S9C**).

The absolute and relative magnitude of deoxygenation did not differ between regions at any dose (**Figure 3G, Suppl Figure S5C**). This dose-dependent decline in sO_2_ occurred independently of changes in total haemoglobin (HbT), which exhibited a nonsignificant upward trend across dosages (**Figure 3F, H; Suppl S5B, D**) but absolute changes in HbT compared to baseline levels or 4 mg/kg did not differ between regions (**Figure 3H, Suppl Figure S5D**).

### Physiological cardiac responses to individual drugs

In the fourth protocol we tested the individual responses to phenylephrine (a selective α□ - adrenergic receptor agonist), isoprenaline (a nonselective β-adrenergic agonist), noradrenaline (a potent agonist at both α□ and β□ adrenergic receptors, with minimal β□ activity), and sodium nitroprusside (to determine whether transient increases in myocardial sO_2_ can be observed, and whether desaturation may result from both hypotension-induced reductions in supply and increased demand, demonstrating that PAI detects both).

As expected, phenylephrine tended to cause a slight initial reduction in heart rate in addition to significantly elevated mean arterial blood pressure (MABP) (∼ 44 mmHg, 95% CI 26.06 to 62.07, P_BP_ = 0.0025) consistent with α-adrenergic increases in total peripheral resistance (**Figure 4A**). Cardiac output in the phenylephrine group trended downward across dosages (**Suppl Figure S6A**). In contrast, isoprenaline tended to cause modest tachycardia (increase of 70 bpm, 95% CI of 13.55 to 148.8, P_HR_=0.046) due to β-adrenergic–mediated positive chronotropy (**Figure 4B**). Isoprenaline led to fluctuations in MABP, and maintained cardiac output across doses (**Figure 4B, Suppl Figure S6B**). Noradrenaline titrations maintained heart rate throughout the experimental recording and led to a trending reduction in cardiac output across dosages which was significant at 6.4 mg/kg (**Figure 4C, Suppl Figure S6C**). Noradrenaline produced a similar elevation in BP as phenylephrine (42 mmHg, 95% CI 30.42-53.47, P_BP_=0.0005) (**Figure 4A, C**). Sodium nitroprusside administration resulted in an immediate and profound reduction in blood pressure (32 mmHg, 95% CI 21.95-41.42, P_BP_=0.0008), but cardiac output did not significantly decrease until 16 mg/kg (**Figure 4D, Suppl Figure S6D**).

**Figure 4.**
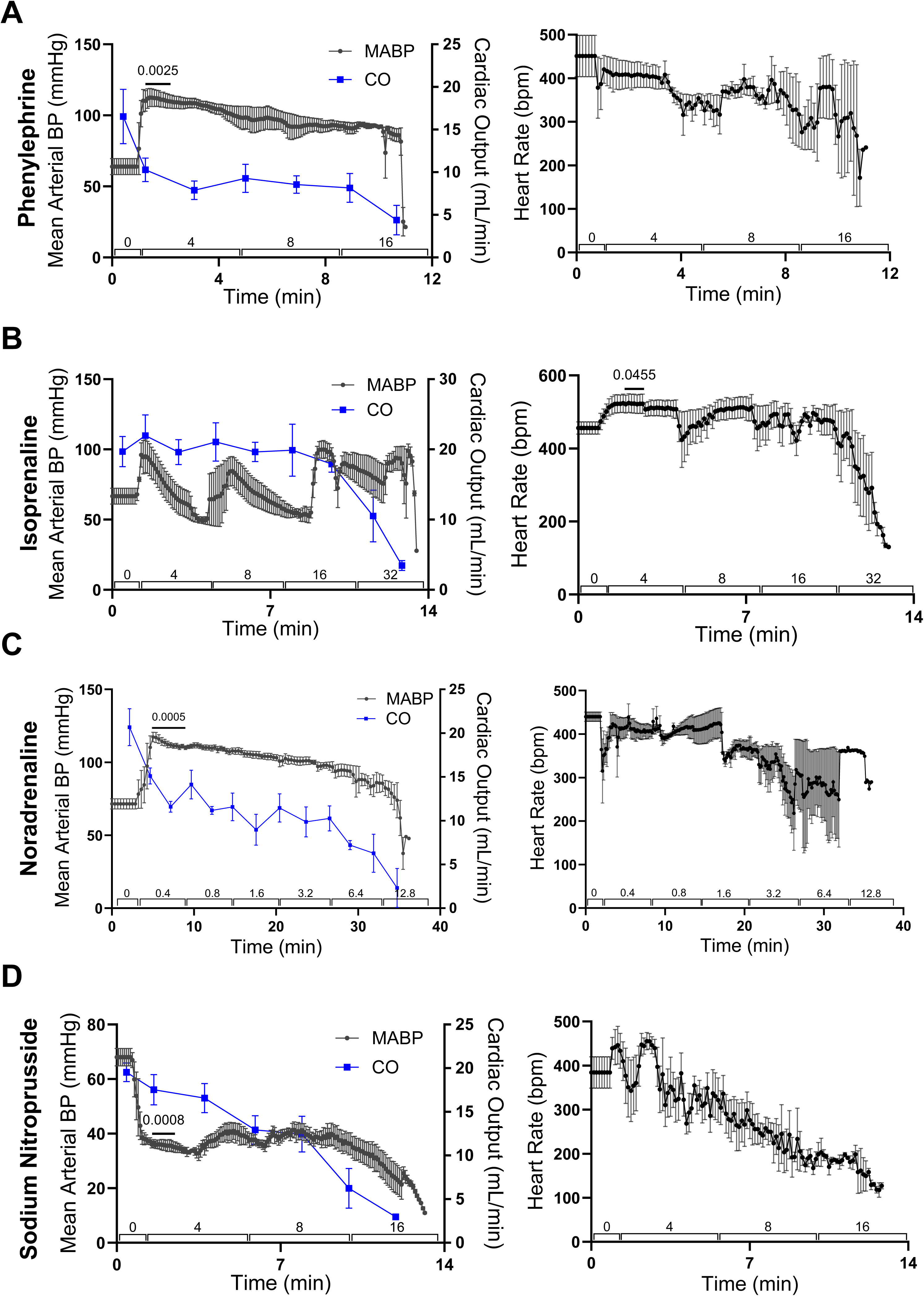
Cardiac physiological responses to individual adrenergic agonists. (**A**) Mean arterial blood pressure (BP) and cardiac output (CO, blue line, right y-axis) changes and heart rate (right) in response to increasing concentrations (doses listed in mg/kg) of phenylephrine. (**B**) Mean arterial blood pressure and cardiac output (blue line, right y-axis) changes and heart (right) rate in response to increasing concentrations (doses listed in mg/kg) of isoprenaline (**C**) Mean arterial blood pressure and cardiac output (blue line, right y-axis) changes and heart rate (right) in response to increasing concentrations (doses listed in mg/kg) of noradrenaline. (**D**) Mean arterial blood pressure and cardiac output (blue line, right y-axis) changes and heart rate in response to increasing concentrations (doses listed in mg/kg) of sodium nitroprusside, n=3 for all panels. Data are displayed as mean±SEM over time. Comparisons between baseline and post-treatment effects were performed using mixed-effects repeated measures 2-way ANOVA followed by Tukey’s post hoc analysis.

With phenylephrine administration, a significant desaturation of the Myo AW and RVOT (mean [95% CI]; Myo AW: -39.95% [-56.78 to -23.12], P=0.0057; RVOT: -29.50 [-33.50 to -25.51], P=0.0017) occurred at 16 mg/kg despite a compensatory increase in HbT at 16 mg/kg (12295 HbT [4143-20447], P= 0.022) (**Figure 5A, Suppl Figure S7A. S8A**). Isoprenaline administration alone elicited a sudden desaturation in the Myo AW and RVOT (mean [95% CI]; Myo AW: -46.97% [-58.77 to -35.16], P=0.019; RVOT: -33.79% [-47.74 to -19.84], P=0.0072) occurring at 32 mg/kg (**Figure 5B, Suppl Figure S7B, S8B** in the context of absent HbT alterations. A gradual myocardial desaturation occurred (mean [95% CI]; Myo AW: -32.02% [-56.80 to -7.24]; trending RVOT: -22.27% [-39.93 to -4.60], P=0.058) with noradrenaline, reaching significance (P=0.049) at a dose of 12.8 mg/kg with no overall changes in HbT across doses (**Figure 5C, Suppl Figure S7C, S8C**). Sodium nitroprusside administration led to an initial increase in Myo AW sO_2_ (3.75%, 95% CI 0.90-6.59, P= 0.022), followed by a substantial desaturation of the Myo AW and RVOT ((mean [95% CI]; Myo AW: -31.51%, [-54.91 to -8.10], P= 0.025; RVOT: -23.24 [-49.31 to 2.82], P=0.025) at a dose of 16 mg/kg (**Figure 5D, Suppl Figure S7D, S8D**).

**Figure 5.**
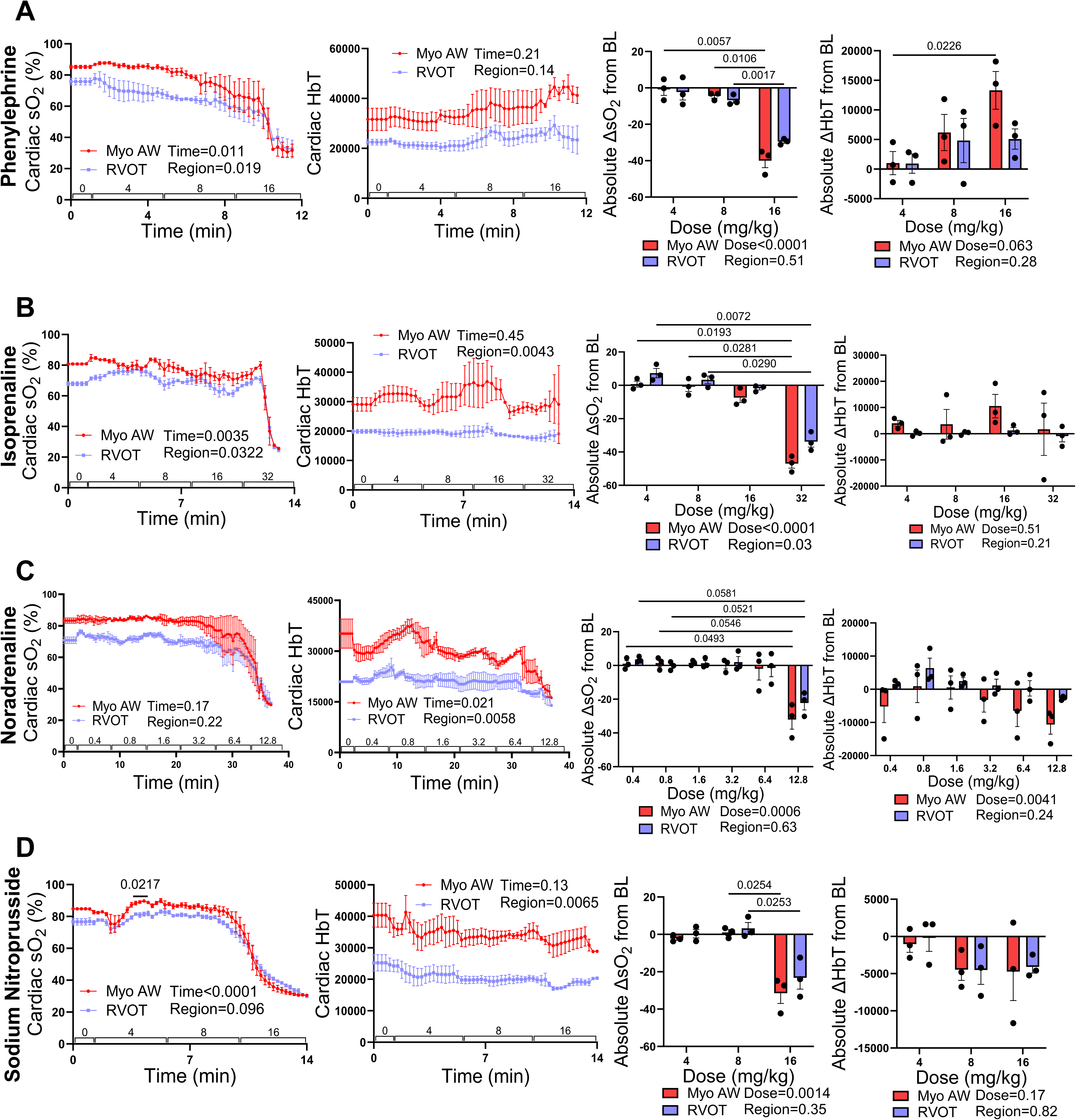
Cardiac oxygenation responses to individual pharmacological challenges. (**A**) Myocardial (Myo AW) and right ventricular outflow tract (RVOT) oxygen saturation (sO_2_, first panel) and total haemoglobin (HbT) concentration (second panel) in response to increasing doses of phenylephrine; quantified absolute changes in Myo AW and RVOT oxygen saturation and haemoglobin concentration across phenylephrine dosages compared to baseline (third and fourth panels respectively). (**B**) Myo AW and RVOT sO_2_ (first panel) and total haemoglobin amounts (HbT, second panel) in response to increasing doses of isoprenaline; quantified absolute changes in Myo AW and RVOT oxygen saturation and HbT across isoprenaline dosages compared to baseline (third and fourth panels). (**C**) Myo AW and RVOT sO_2_ (first panel) and HbT (second panel) in response to increasing doses of noradrenaline; quantified absolute changes in Myo AW and RVOT oxygen saturation and HbT across noradrenaline dosages compared to baseline (third and fourth panels). (**D**) Myo AW and RVOT sO_2_ (first panel) and HbT (second panel) in response to increasing doses of sodium nitroprusside; quantified absolute changes in Myo AW and RVOT oxygen saturation and HbT across sodium nitroprusside dosages compared to baseline (third and fourth panels). n=3 for all panels; Data are displayed as mean±SEM. Comparisons between groups and doses were performed using mixed-effects repeated measures 2-way ANOVA followed by Tukey’s post hoc analysis

Finally, similar to the increases in myocardial perfusion (HbT) observed in protocol three, both phenylephrine and isoprenaline individually elicited trending transient rises in HbT, consistent with metabolic regulation of coronary flow and tone, and phenylephrine-mediated increases in blood pressures (**Figure 5A, B; Suppl Figure S7A-B, S8A-B**). This was not the case with noradrenaline nor sodium nitroprusside, which resulted in a modest trending decrease in HbT compared to baseline (**Figure 5C-D, Figure S7C-D, S8C-D**).

### Myocardial oxygen saturation as an early indicator of hypoxaemia

We assessed whether declines in myocardial sO_2_ preceded changes in cardiac function and electrical alterations, as assessed by left ventricular fractional area change (FAC), and the onset of ECG abnormalities in the settings of hypoxia and drug-induced supply-demand mismatch. During progressive hypoxia, PAI detected a broad range of myocardial sO_2_ values, with FAC remaining stable until myocardial sO_2_ declined to ∼40% (from a baseline of ∼85%) (**Figure 6E**, visually depicted in **Figure 6A-D**). Further, whilst PAI detected changes to myocardial oxygenation between FiO_2_s of 100%, 21% and 10%, GLS (global longitudinal strain) did not reliably detect functional differences between FiO_2_s of between 100%, 21%, and 10% (**Figure 6F**, P_100-10_ = 0.1405). In the setting of hypoxia challenge, reductions in myocardial sO_2_ of up to 20% could be detected prior to the onset of ECG abnormalities (P<0.0001). In 4 of 10 hypoxic mice assessed, ECG abnormalities were not identified (**Figure 6G, Suppl Video 3**).

**Figure 6.**
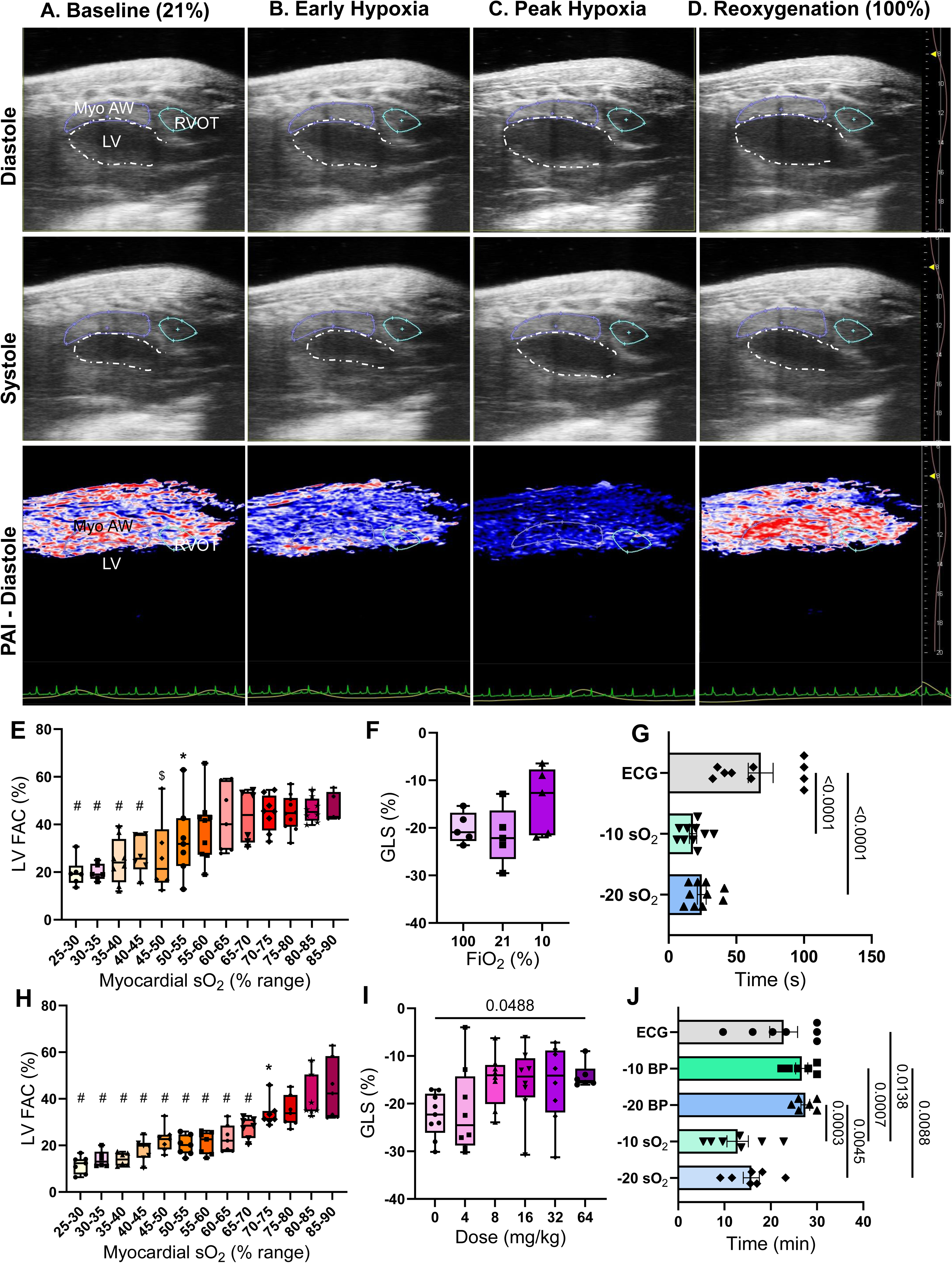
Concurrent B-mode ultrasound and photoacoustic imaging of myocardial oxygenation during hypoxia and reoxygenation and early detection of myocardial hypoxaemia by photoacoustic imaging. Representative parasternal long-axis B-mode ultrasound images with the LV cavity outlined in dashed white acquired at end-diastole (top row) and end-systole (middle row) under baseline normoxia (21% FiO_2_, **A**), early hypoxia (10% at 20 seconds, **B**), peak hypoxia (10% at 60 seconds, **C**), and reoxygenation (100% FiO_2_, **D**). The anterior myocardial wall is delineated in magenta, and the right ventricular outflow tract is outlined in red. Corresponding photoacoustic imaging (PAI) maps acquired continuously and displayed during diastole (bottom row) depict regional myocardial oxygen saturation, where red indicates higher oxygen saturation and blue indicates lower oxygen saturation. Lead I electrocardiogram (ECG) traces are shown in green below PAI images. **(E**) Left ventricular fractional area change (LV FAC, %) plotted across ranges of anterior myocardial wall oxygen saturation in animals exposed to hypoxia challenge, n=6-9. (**F**) Global longitudinal strain (GLS) at steady state FiO_2_s of 100% and 21%, and after 60 s at 10% FiO_2_, n=5. (**G**) Time to reach myocardial anterior wall oxygenation reductions of 10% and 20% from baseline (−10% sO_2_, -20% sO_2_ respectively), and time to reach an electrocardiogram (ECG) abnormality following initiation of recording (100 seconds represents no abnormality), n=10. (**H**) Left ventricular fractional area change (LV FAC, %) plotted across ranges of anterior myocardial wall oxygen saturation (sO_2_) measured during supply-demand mismatch protocols, n=6-7. (**I**) Global longitudinal strain (GLS) at each combined dose of phenylephrine/isoprenaline during supply-demand mismatch, n=8. (**J**) Time to reach 10% and 20% reductions in anterior myocardial wall oxygenation from baseline (−10% sO_2_ and -20% sO_2_, respectively), time to reach reductions in mean arterial blood pressure by 10 and 20 mmHg (−10 BP and -20 BP, respectively), and time to first electrocardiographic (ECG) abnormality following initiation of phenylephrine/isoprenaline administration (30 minutes represents no abnormality), n=6-7. Data are expressed as mean±SEM or mean and IQR, with minimum and maximum values. Comparisons between groups were performed using either a mixed-effects repeated measures 2-way ANOVA followed by the Sidak post hoc test, a one-way ANOVA followed by Dunnett’s multiple Comparisons, or a Mann-Whitney t test. \**p* < 0.05, \*\**p* < 0.01, $*p* < 0.001, #*p* < 0.0001.

In contrast, within our pharmacologically induced myocardial supply-demand mismatch model (**Figure 3A, Suppl Figure S2B**), FAC was reduced at myocardial sO_2_ levels of about ∼70–75%, corresponding to an approximate 15% decrease from baseline (**Figure 6H**). Myocardial systolic function was initially preserved, and FAC was not significantly different between an sO_2_ of 90% (43.90% FAC, 95% CI 32.15–55.65) and 70% (33.72% FAC, 95% CI 28.25–38.98; P_FAC_=0.14) (**Figure 6H, Suppl Video 4**). GLS only significantly declined after the highest dose of cocktail administered, 64 mg/kg (**Figure 6I**). Time to 10% and 20% reductions in myocardial sO_2_ was shorter than time to 10 mmHg and 20 mmHg reduction in blood pressure (P_10BPv10SO2_ =0.0007, P_20BPv20SO2_ =0.0045) (**Figure 6J**). Myocardial sO_2_ reductions of 10% occurred prior to the appearance of ECG abnormalities (P=0.014, **Figure 6J**). In 3 of 7 supply-demand mismatch mice, ECG abnormalities did not occur. The onset of systolic dysfunction in response to drug-induced increased myocardial activity occurred at a higher sO_2_ compared to that of our hypoxia challenges, and may reflect direct myocardial effects of the administered agents, as well as a more gradual temporal decline in oxygenation following prolonged periods of mild myocardial stress at lower pharmacological doses. Nonetheless, during the post-dose monitoring period after administration of 16 mg/kg, persistent myocardial hypoxemia was occasionally observed without accompanying electrical or functional abnormalities (**Suppl Video 4**).

## DISCUSSION

Within the “ischaemic cascade”, cardiac hypoxia or ischaemia precedes electrical or mechanical manifestations, necessitating strategies to detect myocardial oxygenation to prevent ensuing myocardial tissue injury and functional decline. Here we demonstrate PAI’s capability to measure oxygenation within the myocardium, RVOT, pulmonary artery and aorta and monitor real-time responses to hypoxic and pharmacological insults.

Photoacoustic imaging enabled detectable changes in sO_2_ of the above regions in response to moderate reductions in oxygen delivery from FiO_2_s of 100% to 21%. However, to obtain a higher resolution image we then attempted to evaluate EKV (Electrocardiogram-gated Kilohertz Visualisation). EKV PAI demonstrated the ability to resolve sO_2_ differences between the myocardium and RVOT under FiO_2_ conditions of 100% and 21% but, in this application, offered no clear advantage over conventional non-gated photoacoustic imaging. This may be attributed to the averaging of measurements, which could introduce noise as subtle shifts in cardiac position occur between cycles. This further underscored the robust temporal-spatial resolution of standard photoacoustic imaging, which effectively resolved sO_2_ differences without the need for motion gating. Therefore, standard PAI allowed for feasibility of subsequent experiments, permitting continuous monitoring of oxygenation. PAI aortic sO_2_ measurements closely approximated peripheral arterial pulse oximetry within physiological sO_2_ ranges; at lower sO_2_s, however, PAI produced more reliable measurements than pulse oximetry, which frequently lost signal fidelity.

While oxygen delivery and extraction are routinely measured, the study herein is the first of its kind to assess real-time responses of myocardial oxygenation in response to hypoxic episodes. In response to an acute prominent hypoxic challenge (FiO_2_ 10%), the aortic sO_2_ and pulmonary artery sO_2_ declined similarly, but to our surprise, myocardial oxygenation demonstrated a more rapid and pronounced decline than the RVOT. The reduction of myocardium sO_2_ to levels lower than that of systemic venous oxygen saturation (measured clinically by mixed venous blood gas and experimentally by RVOT sO_2_ in our model) may identify an adaptive increase in myocardial oxygen utilization, or highlight the limited reserve of the heart which may underlie myocardial susceptibility to injury from various stressors during the perioperative period ^32, 33^. Although our hypoxia-reoxygenation studies were executed to provide proof of principle supporting PAI’s utility, this model of rapid hypoxia represents the clinical scenario of a loss of airway or asphyxiation. The nadir myocardial and great vessel sO_2_ signals of 25-30% were consistent with physiologically plausible residual hemoglobin oxygen saturation.

Importantly, in conditions of severe hypoxia, we found that PAI identified marked reductions in myocardial sO_2_ well before ECG and functional abnormalities were evident, and frequently, in the complete absence of ECG changes. During acute hypoxia, autoregulation of coronary vasculature maintains flow to preserve oxygen delivery ^34^; as such, the observed reductions in sO_2_ in response to hypoxia likely reflect increased oxygen extraction rather than decreased perfusion.

To model supply-demand mismatch, we administered increasing doses of phenylephrine and isoprenaline (delivered concurrently in a two to one ratio), simulating a clinical scenario of vasopressor and inotropic support during shock that may ultimately lead to cardiovascular collapse. This approach produced increased cardiac demand via heightened afterload and workload, resulting in progressive dose-dependent reductions in myocardial and RVOT sO_2_. This reduction in sO_2_ likely reflects increased myocardial oxygen extraction and utilization to meet heightened metabolic demand ^31^, occurring alongside a trend toward increased myocardial perfusion during cocktail titration, which reached significance with phenylephrine alone ^28, 30^. Myocardial desaturation became profound at intermediate doses while cardiac performance was initially maintained and reduced only at high doses. Notably, blood samples showed elevated cardiac troponin and lactate, demonstrating biochemical evidence of myocardial injury.

When compared to a hypoxia challenge of FiO_2_ 10%, the drug-induced onset of systolic dysfunction occurred at a relatively higher myocardial sO_2_ (i.e. FAC decreased after a 15-20% sO_2_ reduction in response to demand ischaemia as opposed to a 45-50% sO_2_ reduction in hypoxia) due to various factors. This difference may be due to increased afterload from phenylephrine, combined with increased oxygen requirements secondary to the phenylephrine and isoprenaline, as well as the direct effect of high doses of adrenergic agonists on myocardial receptors. The gradual increase in phenylephrine and isoprenaline titrations as opposed to the immediate reduction in FiO_2_ led to prolonged periods of cardiac stress and potential decrease in cardiac reserve. In the pharmacologically induced supply-demand mismatch, myocardial sO_2_ decreased by 20% before blood pressure declined by 10 mmHg, and ECG aberrations appeared once myocardial sO_2_ approached a ∼20% reduction from baseline values. However, within the hypoxia challenge, ECG aberrations occurred well after this 20% reduction in myocardial sO_2_ or did not occur at all. Collectively, these findings suggest that photoacoustic imaging detects clinically meaningful declines in myocardial oxygen saturation preceding both functional impairment and electrical abnormalities, highlighting its potential for early identification of impending cardiovascular collapse.

In addition to measuring the cardiac response to a combination of phenylephrine and isoprenaline, we sought to characterise the myocardial response to these medications individually. Given that phenylephrine and noradrenaline have similar clinical indications, noradrenaline was also assessed due to its more favourable myocardial oxygenation profile secondary to its modest β-agonism. We found that phenylephrine appeared to elicit myocardial desaturation more rapidly and at a lower dose than the combined phenylephrine and isoprenaline used previously, both of which elicited desaturation at a relatively lower dose than noradrenaline. Delayed desaturation associated with noradrenaline, an endogenous catecholamine, may be due to its evolved ideal balance of α□, β□, and β□ activation to support haemodynamics. These findings may reflect the distinct receptor profiles of the agents. Phenylephrine, as a pure α□- agonist, produces a more abrupt increase in systemic vascular resistance without β□-mediated support of cardiac output. The resulting combination of heightened afterload and the tendency for even modest reflex bradycardia could accelerate the onset of myocardial desaturation, even at comparatively lower doses. The addition of isoprenaline provided β□-mediated support of cardiac output, coupled with modest β□-mediated coronary vasodilation, delaying the onset of desaturation. Sodium nitroprusside produced a transient increase in myocardial sO_2_ accompanied by profound hypotension, but ultimately led to myocardial desaturation at a dose comparable to phenylephrine, despite opposing systemic blood pressure effects; this demonstrates that PAI detects desaturation arising from both supply and demand failure.

Although PAI has been used in clinical research it has mostly been adopted to measure oxygenation or distinguish between various soft tissue types ^35, 36^. PAI could be useful in several clinical scenarios as it offers non-invasive, dynamic, and spatial monitoring of myocardial and great vessel oxygenation (albeit with reduced penetration depth). PAI may thus inform accurate titration of FiO_2_, pharmacological agents, blood products, and ventilator settings to improve haemodynamic control, particularly in patients with poor reserve, in addition to providing an earlier warning sign of impending haemodynamic collapse. The ability to measure oxygenation within the aorta offers a more central measure of continuous oxygenation than pulse oximetry, and may offer more reliable measurements during poor peripheral perfusion, and evade delays in identifying hypoxaemia, which can range from 20-120 s ^19–22^. In veno-arterial extracorporeal membrane oxygenation (ECMO), locating the watershed line along the aorta to inform early initiation of veno-arterial-venous ECMO could prevent cardiac arrest and stroke ^37, 38^. Finally, the ability to trend pulmonary arterial oxygenation may inform the management of right heart failure in patients with pulmonary hypertension.

Given the ubiquity of ultrasound, PAI could be readily integrated into the existing monitoring platforms as a standard of care. However, depth of penetration remains an issue for adult humans; PAI may therefore offer a practical first-line option for paediatric assessments and for adult epicardial measurements during cardiac surgery and could become increasingly feasible with the development of transoesophageal probes. Finally, the development of more powerful lasers and more sensitive ultrasound transducers is underway, which would improve depth of penetration.

### Limitations

This study has certain limitations. Photoacoustic imaging is subject to depth-dependent signal attenuation, subtly impacting precision and precluding intra-ventricular blood sO_2_ measurements. Differences in optical attenuation whereby a loss in signal intensity as light traverses various tissues at increased depths can affect sO_2_ measurements ^39^. This can be circumvented with time gain compensation, perhaps explaining why pulmonary artery and right ventricular outflow tract oxygenation did not significantly differ despite being measured from distinct views and depths. The small size of murine models limited the integration of comprehensive physiological measurements within a single preparation. Coronary blood flow could not be reliably assessed due to the small calibre and intramyocardial location of murine coronary vessels, and the current Vevo LAZR-X platform does not permit separation of arterial and venous compartments required for oxygen extraction calculations. Direct blood gas sampling was also not feasible due to limited circulating volume, and pulse oximetry was used as a surrogate for arterial oxygenation. Future studies in larger animal models may enable comprehensive physiological characterisation, including coronary flow measurement and blood sampling. Finally, as photoacoustic imaging measures the ratio of oxygenated and deoxygenated haemoglobin, it likely would not detect impairments of intracellular oxygenation.

In conclusion, photoacoustic imaging identifies the early myocardial oxygen imbalance that arises during hypoxic and pharmacologic stress, enabling detection prior to electrical and functional deterioration. We show that the myocardium is acutely vulnerable to oxygen imbalance, with rapid desaturation reflecting increased oxygen extraction in the face of limited reserve. Furthermore, we identified differential myocardial tolerance to various pressors, with noradrenaline delaying myocardial hypoxaemia. This study propounds the potential for PAI to be integrated into the existing monitoring armamentarium, and showcases the value of PAI in detecting early shifts in cardiovascular oxygenation before conventional indicators, which may in turn enable timelier, targeted, and effective intervention strategies.

## Author contributions

Conceptualization: JV, RN, SLB, MH, SD

Methodology, investigation and experimentation: JV, RN, IK

Data analysis and figure generation: JV, RN, SLB

Supervision and provision of equipment: SLB, RN, STD, KFM

Writing - original draft: JV, RN

Writing - review and editing: JV, RN, SD, MH, SLB, KFM

Funding acquisition: SLB, STD, JV

All authors reviewed and approved the final manuscript.

## SUPPLEMENTAL FIGURE LEGENDS

**Figure S1.** (**A**) Experimental setup with mouse instrumented with a femoral cannula attached to a syringe with drug cocktail; the ultrasound transducer is equipped with a laser jacket containing two laser inserts, the transducer is positioned parasternally. The mouse and laser are contained within a VevoLAZR barrier box. (**B**) Experimental group design layout

**Figure S2.** (**A**) Experimental timeline of hypoxia challenges. FiO_2_ = fraction of inspired oxygen, min. = minute. (**B**) Experimental timeline of supply-demand mismatch protocol and phenylephrine-isoprenaline cocktail administration. GLS = global longitudinal strain, CO = cardiac output. (**C**) Analysis and comparison of electrocardiogram-kilohertz visualization (EKV) mode and standard photoacoustic imaging (Std PAI) mode derived myocardial sO_2_ values captured at end-systole and end-diastole, n=8. (**D**) Measures of variance (standard deviation and coefficient of variation) between EKV and Std PAI modes during steady state baseline recordings of myocardial sO_2_, n=5. (**E**) Comparison of right ventricular outflow tract (RVOT, obtained in a long axis view) and pulmonary artery, (Pulm Art, visualized in a short axis view) sO_2_ values adjusted with time gain compensation to account for angle and depth differences across FiO_2_ conditions of 100 and 21%, n=6. Data are expressed as mean and IQR, with minimum and maximum values. Data was analyzed with 2-way ANOVA followed by Tukey’s post hoc test, or by paired t-test.

**Figure S3.** (**A**) Oxygen saturation (left) and haemoglobin concentration (right) over time across the transition from steady state FiO_2_s of 100% (baseline) to 21% of the myocardial anterior wall (Myo AW) and right ventricular outflow tract (RVOT), n=9. (**B**) Oxygenation (left) and haemoglobin concentration (right) over time across the transition from steady state FiO_2_s of 100% (baseline) to 21% of the aorta and pulmonary artery (Pulm Art), n=5. (**C**) Oxygen saturation (left) and haemoglobin concentration (right) of cardiac regions (including Myo AW and RVOT) during the transition from FiO_2_ 21 to 0 (no further decline in myocardial sO_2_ observed following these recordings), n=5. (**D**) Oxygen saturation (left) and haemoglobin concentration (right) of great vessels (including aorta and pulmonary artery) during the transition from FiO_2_ 21 to 0, n=4. Data are expressed as mean±SEM. Comparisons between groups were performed using mixed effects repeated measures 2-way ANOVA with Tukey’s post hoc analysis.

**Figure S4.** (**A**) Relative changes in oxygenation (sO_2_) between the Myo AW and RVOT chamber in response to FiO_2_ transitions from 21 to 10 to 100% (above); Relative changes in haemoglobin concentration (HbT) between the Myo AW and RVOT chamber in response to FiO_2_ transitions from 21 to 10 to 100% (below), n=11. (**B**) Relative changes in oxygenation (sO_2_) between the aorta and pulmonary artery in response to FiO_2_ transitions from 21 to 10 to 100% (above); Relative changes in haemoglobin concentration (HbT) between the aorta and pulmonary artery in response to FiO_2_ transitions from 21 to 10 to 100% (below), n=4. (**C**) Relative changes in oxygenation (left) and haemoglobin concentration (right) from baseline in the Myo AW and RVOT in response to FiO_2_ transitions of 100% to 21% and 21% to 10%. (**D**) Relative changes in oxygenation (left) and haemoglobin concentration (right) from baseline in the aorta and pulmonary artery in response to FiO_2_ transitions of 100% to 21% and 21% to 10%. Data are expressed as mean±SEM. Comparisons between groups were performed using either a mixed-effects repeated measures 2-way ANOVA followed by Tukey’s post hoc test, \**p* < 0.05, \*\**p* < 0.01, $*p* < 0.001, #*p* < 0.0001.

**Figure S5.** (**A**) Normalised alterations in oxygenation of the Myo AW and RVOT in response to increasing phenylephrine and isoprenaline cocktail inducing progressive supply-demand mismatch. (**B**) Normalised changes in total haemoglobin of the Myo AW and RVOT in response to increasing phenylephrine and isoprenaline cocktail inducing progressive supply-demand mismatch. (**C**) Relative changes in oxygenation of the Myo AW and RVOT compared to baseline in response to increasing dosages of phenylephrine and isoprenaline cocktail. (**D**) Relative changes in haemoglobin concentration in the Myo AW and RVOT compared to baseline in response to increasing dosages. n=4-8 for all panels, Data are expressed as mean±SEM. Comparisons between groups were performed using either a mixed-effects repeated measures 2-way ANOVA followed by Tukey’s post hoc test, *p < 0.05, **p < 0.01, $p < 0.001, #p < 0.0001.

**Figure S6.** Cardiac output measured by echocardiography across intravenously administered doses of individual adrenergic or vasoactive agents: (**A**) phenylephrine, (**B**) isoprenaline, (**C**) noradrenaline, and (**D**) sodium nitroprusside. Data are displayed as mean and IQR. Comparisons between groups were performed using repeated measures one-way ANOVA followed by Tukey’s multiple comparisons test.

**Figure S7.** Normalised myocardial (Myo AW) and right ventricular outflow tract (RVOT) oxygenation (left panel) and total haemoglobin concentration (middle panel), and heart rate in response to increasing concentrations (doses listed in mg/kg) of (**A**) phenylephrine, (**B**) isoprenaline, (**C**) noradrenaline, and (D) sodium nitroprusside. n=3 for all panels; Data are displayed as mean±SEM over time. Comparisons between groups were performed using mixed-effects repeated measures 2-way ANOVA followed by Tukey’s post hoc analysis.

**Figure S8.** (**A**) Normalised changes in Myo AW and RVOT oxygenation (left) and haemoglobin concentration (right) in response to increasing doses of phenylephrine. (**B**) Relative changes in Myo AW and RVOT oxygenation (left) and total haemoglobin (right) in response to increasing doses of isoprenaline. (**C**) Normalised changes in Myo AW and RVOT oxygenation (left) and haemoglobin concentration (right) in response to increasing doses of noradrenaline. (**D**) Normalised changes in Myo AW and RVOT oxygenation (left) and haemoglobin concentration (right) in response to increasing doses of sodium nitroprusside. n=3 for all panels; Data are displayed as mean±SEM and analyzed with repeated measures 2-way ANOVA followed by Tukey’s post hoc analysis.

**Figure S9.** (**A**) Representative individual tracings following sequential FiO_2_s of 21% (baseline), 10% (60s) and 100% (90s) of the myocardial anterior wall (green) and right ventricular outflow tract (RVOT, blue) (**B**) Representative individual tracings following sequential FiO_2_s of 21% (baseline), 10% (60s) and 100% (90s) of the aorta (green) and pulmonary artery (red). (**C**) Representative individual traces of the myocardial anteroseptal wall and RVOT during critical desaturation periods resulting from pharmacologically induced (phenylephrine and isoprenaline-induced) supply-demand mismatch; the myocardial anterior wall is depicted in green and the RVOT is depicted in orange for both recordings. Images were generated and obtained with Vevo Lab 5.8.0.

**Video V1.** Representative recording of the myocardial anterior wall and right ventricular outflow tract oxygen saturation in real time resulting in sudden hypoxaemia after administration of 16 mg/kg phenylephrine-isoprenaline cocktail.

**Video V2.** Second representative recording of the myocardial anterior wall and right ventricular outflow tract oxygen saturation leading to gradual and progressive hypoxaemia after administration of 32 mg/kg of phenylephrine-isoprenaline cocktail.

**Video V3.** Representative recording of the myocardial anterior wall and right ventricular outflow tract following FiO_2_s of 21% (baseline), 10% and 100%. FiO_2_ changed from 21% to 10% at approximately 6.27s, and from 10% to 100% at approximately 36.27s in the absence of ECG abnormalities.

**Video V4.** Representative recording of myocardial and right ventricular tract oxygen saturation during persistent cardiac hypoxemia in response to combined drug-induced supply-demand mismatch (32 mg/kg of phenylephrine-isoprenaline) in the absence of ECG abnormalities

