## Supplemental Figures for "Real-Time Assessment of Murine Cardiac Oxygenation Using Photoacoustic Imaging"

### Slide 1
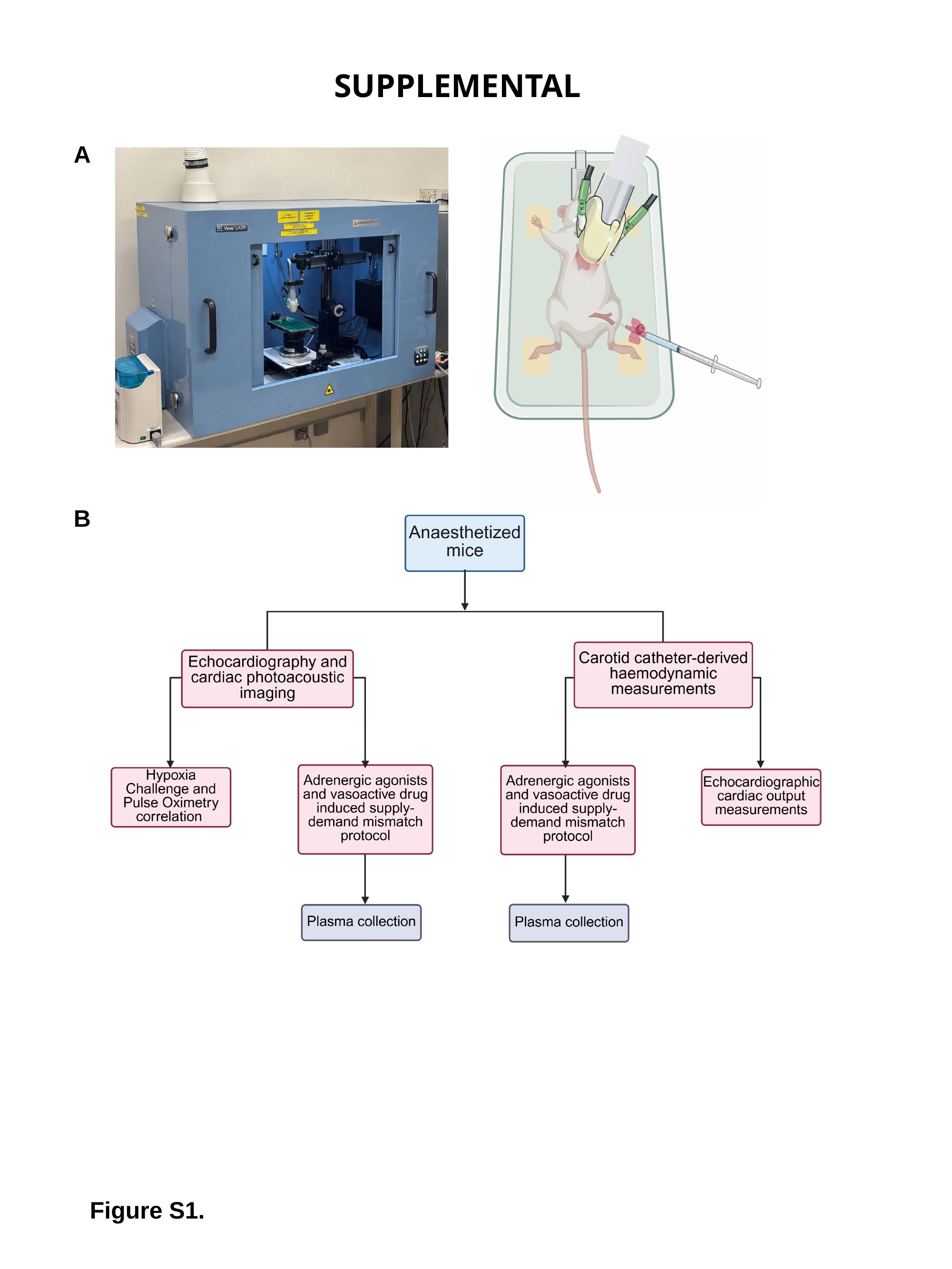

SUPPLEMENTAL
A
B
Figure S1.

### Slide 2
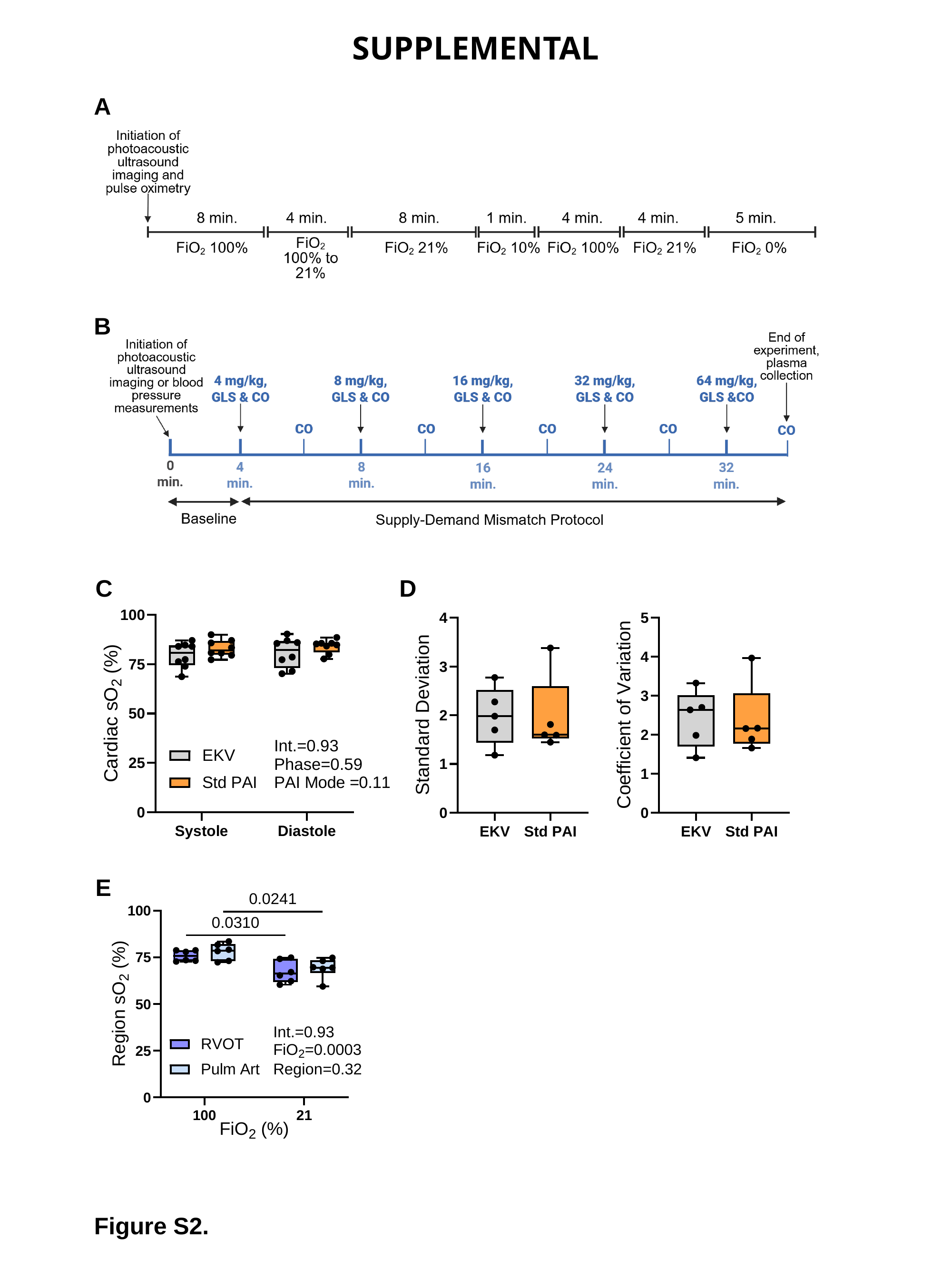

SUPPLEMENTAL
A
B
C
D
E
Figure S2.

### Slide 3
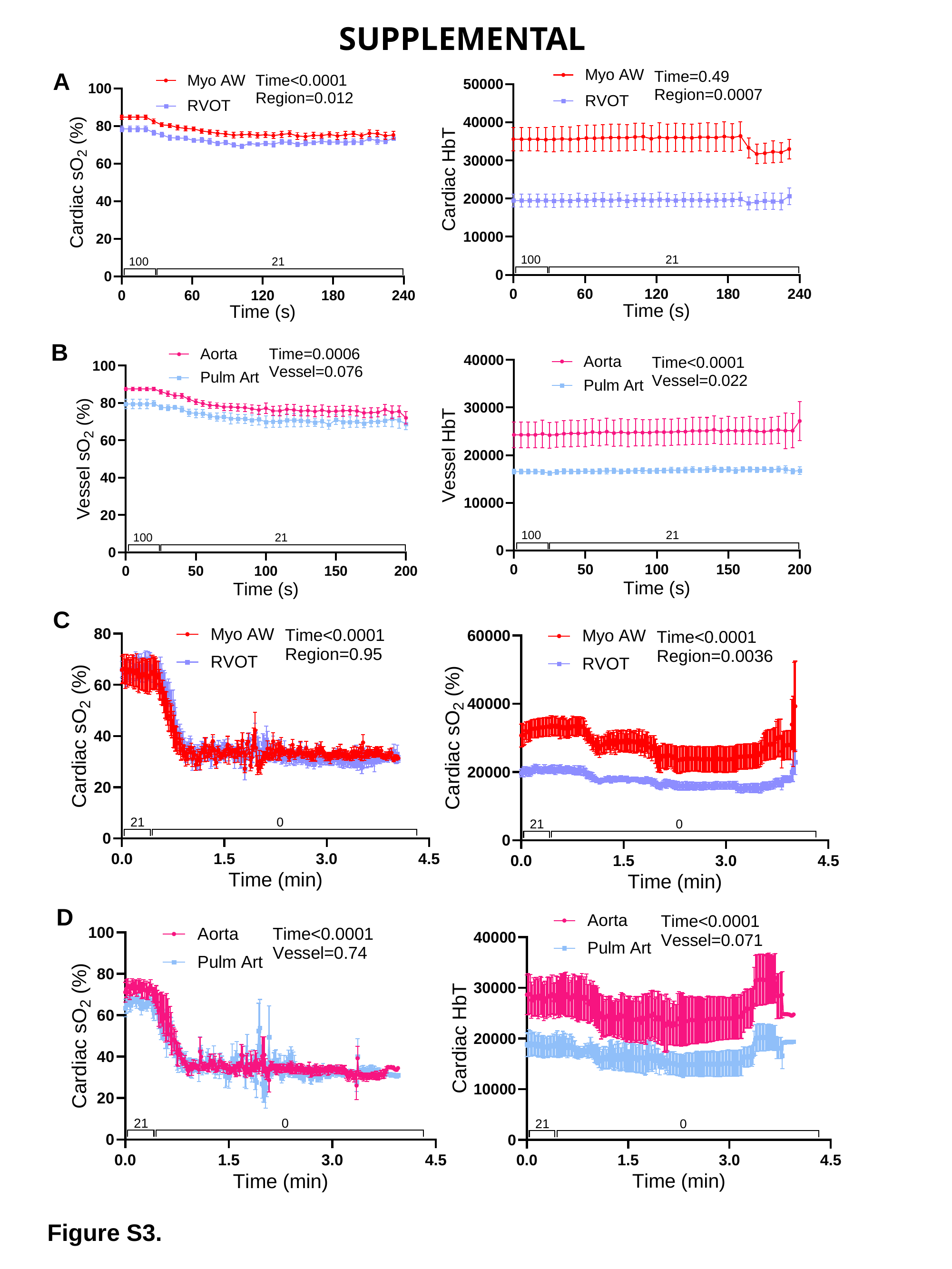

SUPPLEMENTAL
A
B
C
D
Figure S3.

### Slide 4
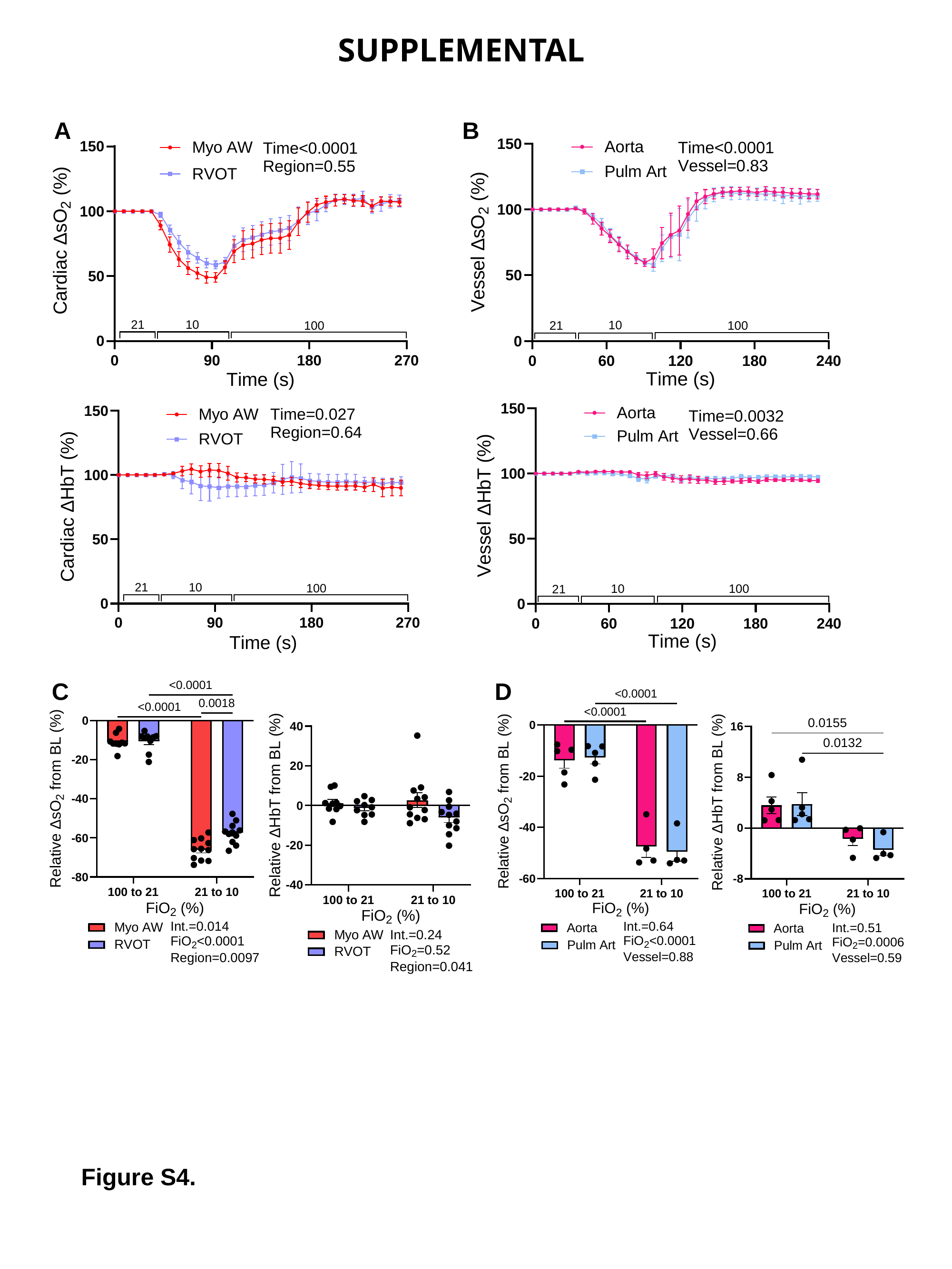

SUPPLEMENTAL
A
B
C
D
Figure S4.

### Slide 5
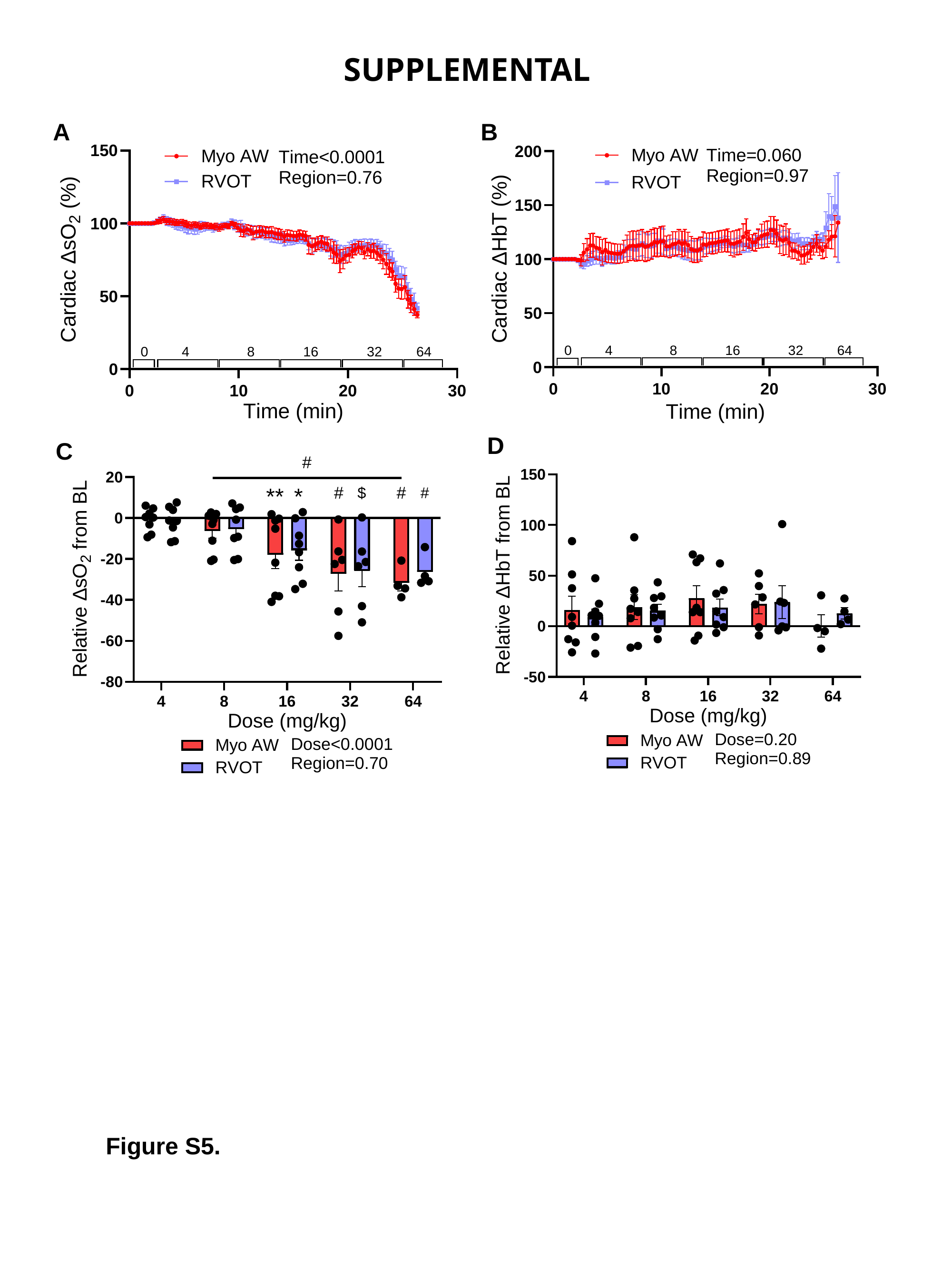

SUPPLEMENTAL
B
A
D
C
Figure S5.

### Slide 6
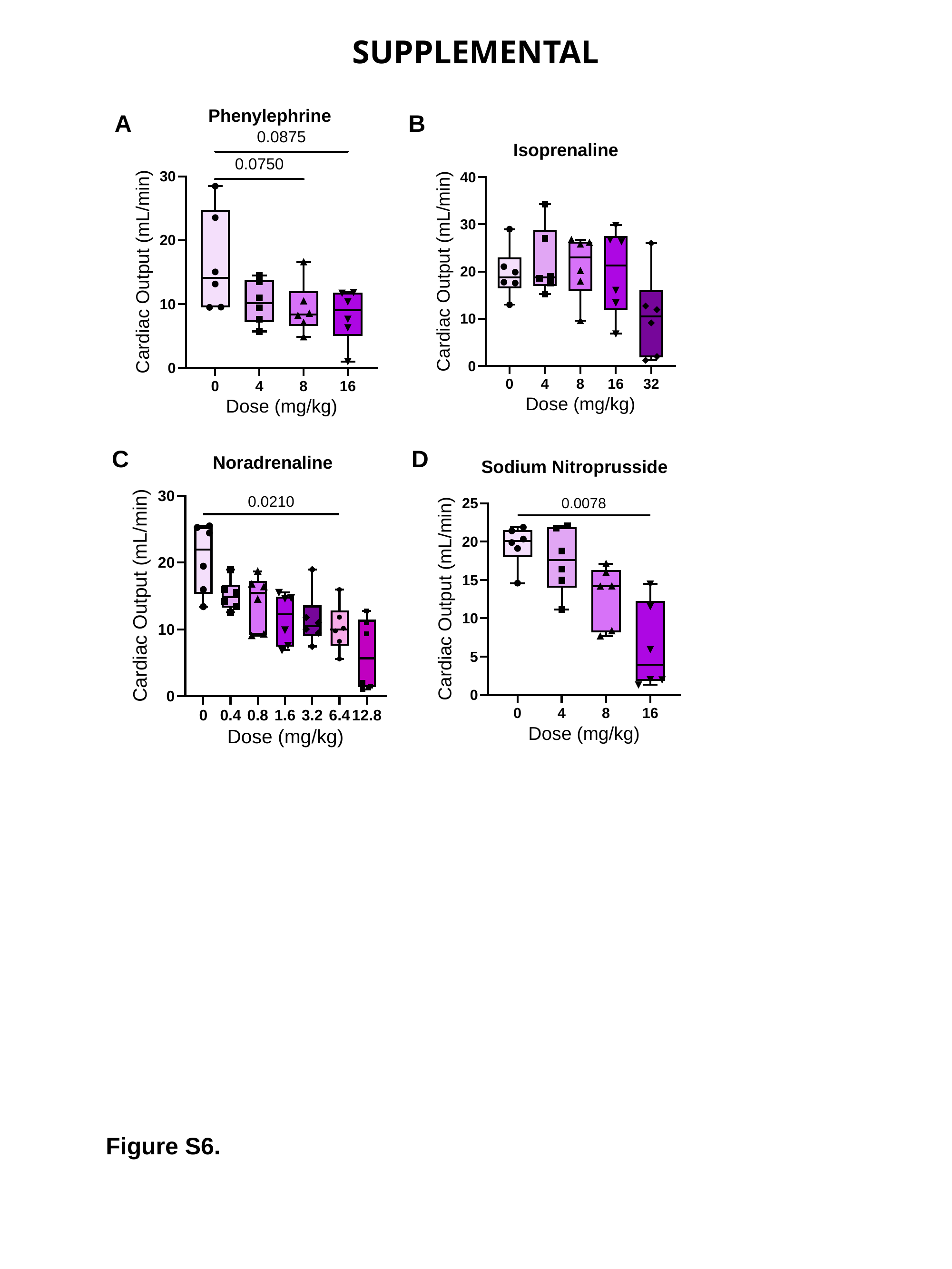

SUPPLEMENTAL
Phenylephrine
A
B
Isoprenaline
C
D
Noradrenaline
Sodium Nitroprusside
Figure S6.

### Slide 7
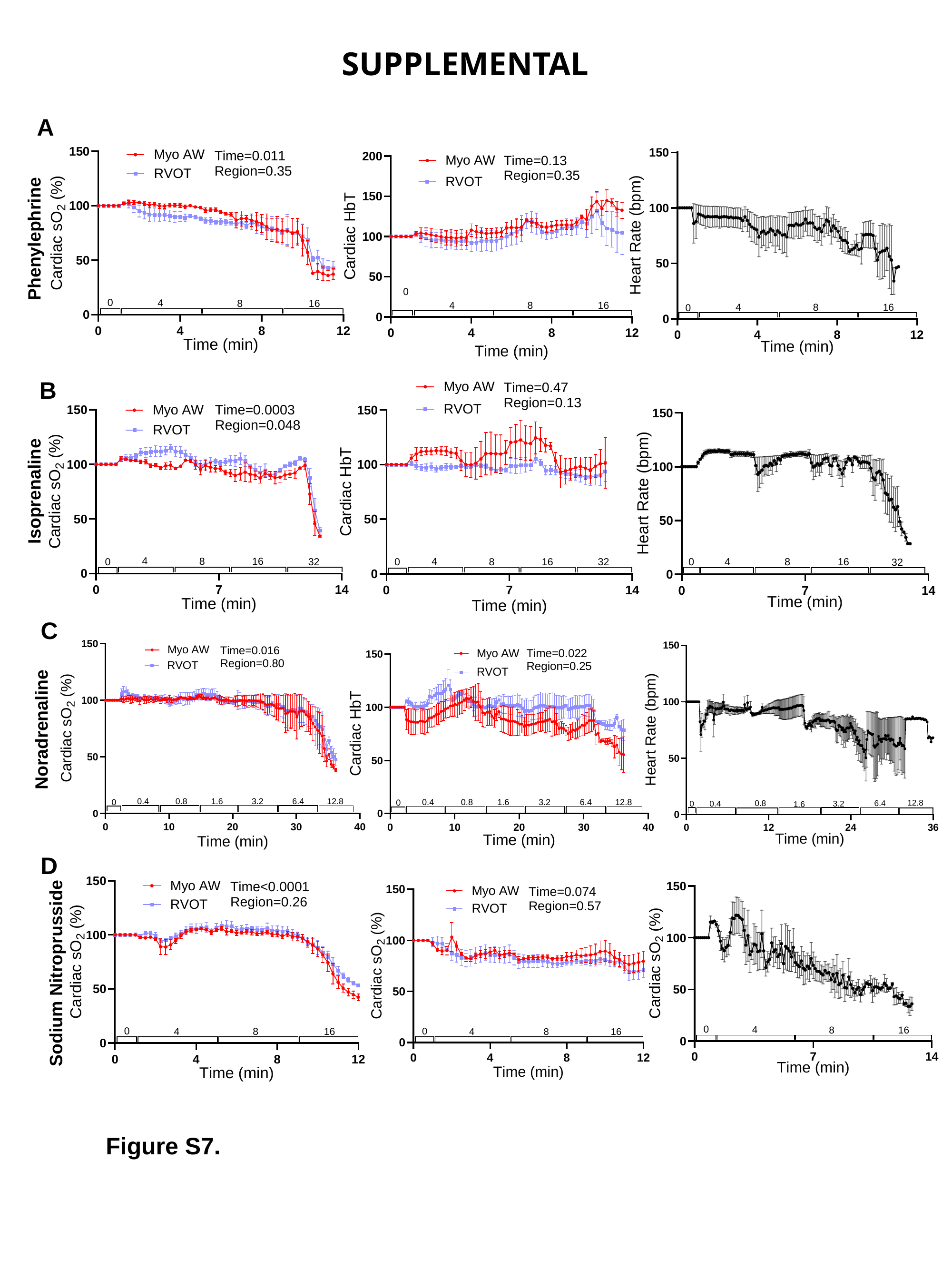

SUPPLEMENTAL
A
Phenylephrine
B
Isoprenaline
C
Noradrenaline
D
Sodium Nitroprusside
Figure S7.

### Slide 8
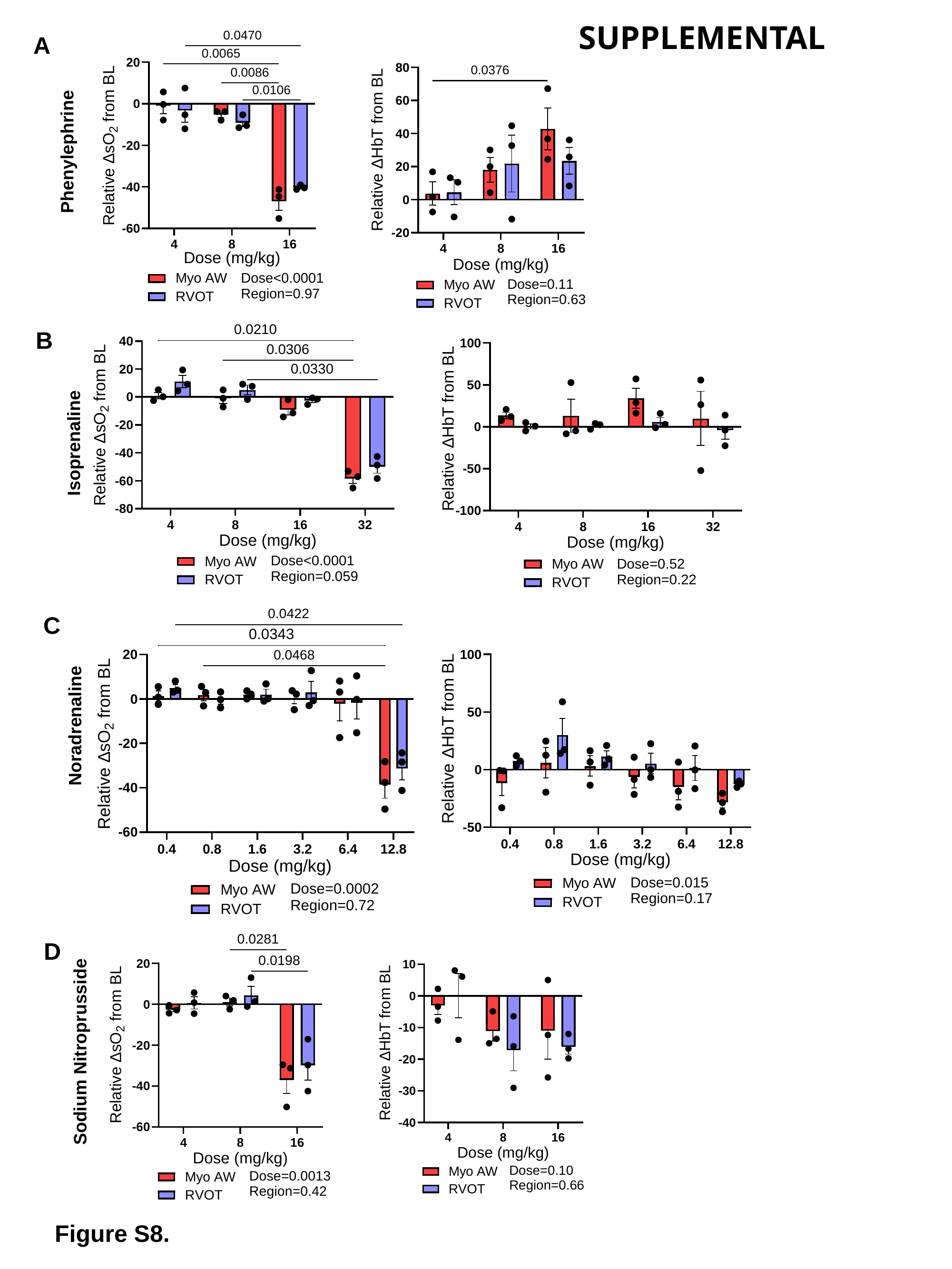

SUPPLEMENTAL
A
Phenylephrine
B
Isoprenaline
C
Noradrenaline
D
Sodium Nitroprusside
Figure S8.

### Slide 9
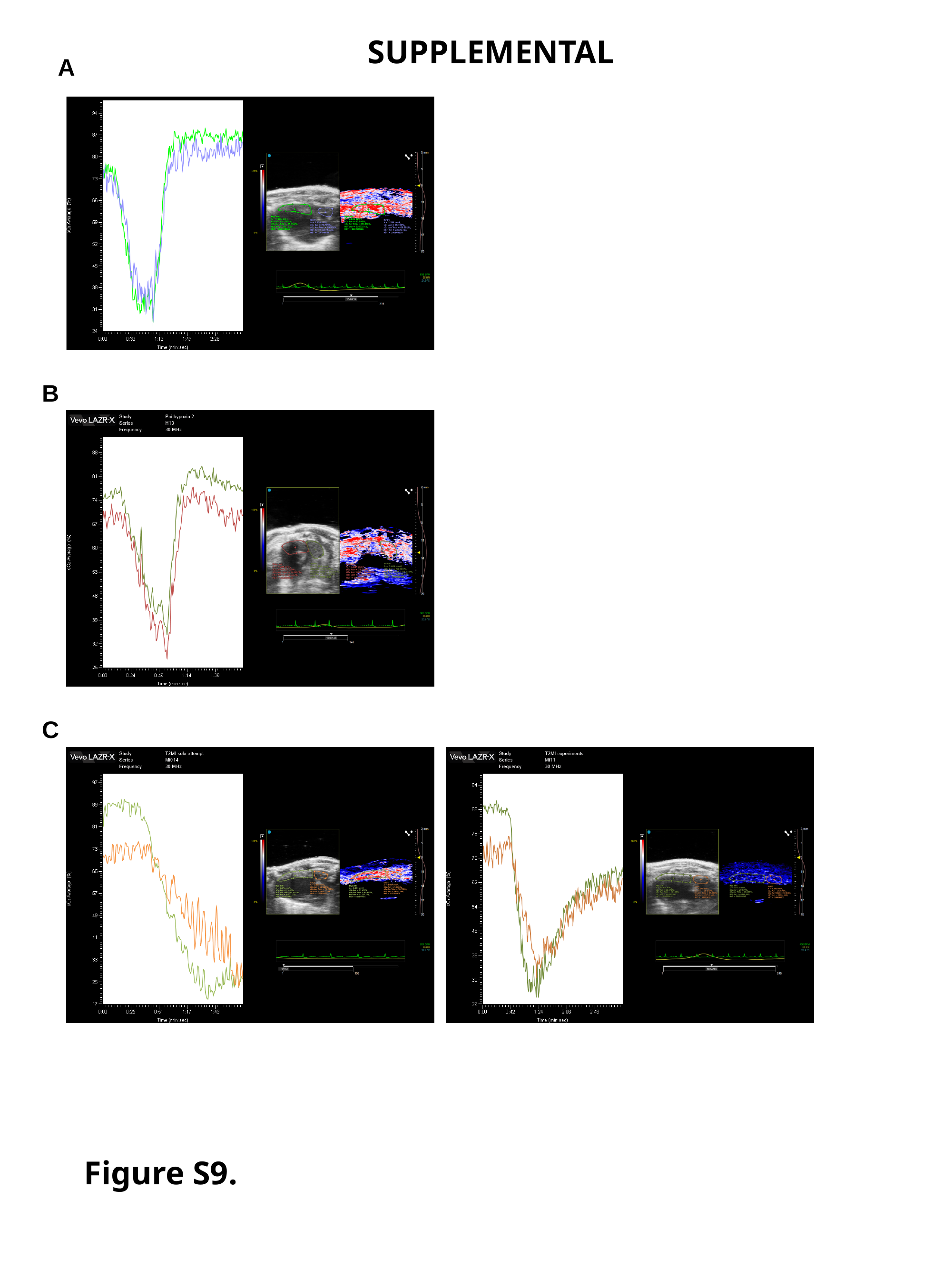

SUPPLEMENTAL
A
B
C
Figure S9.
